# Glutathione in the nucleus accumbens regulates motivation to exert reward-incentivized effort

**DOI:** 10.1101/2022.02.14.480343

**Authors:** Ioannis Zalachoras, Eva Ramos-Fernández, Fiona Hollis, Laura Trovò, João Rodrigues, Alina Strasser, Olivia Zanoletti, Pascal Steiner, Nicolas Preitner, Lijing Xin, Simone Astori, Carmen Sandi

**Affiliations:** Laboratory of Behavioral Genetics (LGC), Brain Mind Institute, École Polytechnique Fédérale de Lausanne (EPFL), Lausanne, Switzerland; Nestlé Institute of Health Sciences, Nestlé Research, Société des Produits Nestlé S.A., Vers-chez-les-Blanc, 1000 Lausanne 26, Switzerland; Animal Imaging and Technology Core (AIT), Center for Biomedical Imaging (CIBM), Ecole Polytechnique Fédérale de Lausanne (EPFL), Lausanne, Switzerland

## Abstract

Emerging evidence is implicating mitochondrial function and metabolism in the nucleus accumbens in motivated performance. However, the brain is vulnerable to excessive oxidative insults resulting from neurometabolic processes and whether antioxidant levels in the nucleus accumbens contribute to motivated performance is not known. Here, we identify a critical role for glutathione (GSH), the most important endogenous antioxidant in the brain, in motivation. Using proton magnetic resonance spectroscopy (^1^H-MRS) at ultra-high field in both clinical and preclinical populations, we establish that higher accumbal GSH levels are highly predictive of better, and particularly steady performance over time in effort-related tasks. Causality was established in preclinical in vivo experiments that, first, showed that down-regulating GSH levels through micro-injections of the GSH synthesis inhibitor buthionine sulfoximine in the nucleus accumbens impaired effort-based reward-incentivized performance. In addition, systemic treatment with the GSH precursor N-acetyl-cysteine (NAC) increased accumbal GSH levels and led to improved performance, potentially mediated by a cell-type specific shift in glutamatergic inputs to accumbal medium spiny neurons. Our data indicate a close association between accumbal GSH levels and individual’s capacity to exert reward-incentivized effort over time. They also suggest that improvement of accumbal antioxidant function may be a feasible approach to boost motivation.

## Introduction

Motivation facilitates overcoming the cost of effortful actions to attain desired outcomes^1^ and is key to achievement and well-being^2–4^. Importantly, there are substantial differences in motivated behavior among healthy individuals, manifested as variations in the engagement in effortful activities and in differences in brain activity^5,6^. Motivational deficits - such as apathy, anhedonia or anergia- are prevalent in many brain pathologies^7–10^. Therefore, unveiling the neurobiological mechanisms underlying individual differences in motivation can help develop novel interventions to boost effortful performance.

A great deal of work in both rodents and humans highlights the nucleus accumbens (NuAc) - a main component of the ventral striatum- as a critical node of the brain’s reward and motivation circuitries^9,11–14^. Indeed, incentives energize effortful behavior^6,15^ through the recruitment of the NuAc^14,16–21^. Moreover, alterations in NuAc function are implicated in psychopathological conditions, that involve motivational deficits, such as depression^22–25^.

Emerging evidence underscores a key role for accumbal mitochondrial function and metabolism in the regulation of motivated behavior^26–29^ and vulnerability to develop stress-induced depressive-like behaviors^30–32^. This is in line with the established relevance of cellular and tissue energy metabolism to maintain healthy brain function^33,34^, and goes beyond to posit a link between variation in the levels of specific accumbal metabolites and motivated behavior. However, knowledge regarding the specific accumbal metabolites involved in effortful performance is still scarce.

Recent work in humans using proton magnetic resonance spectroscopy (^1^H-MRS) focusing on the components of the glutamate/GABA-glutamine cycle has shown that glutamine levels, and particularly an increased glutamine-to-glutamate ratio in the NuAc predict both, a higher performance related to task endurance and a lower effort perception in a physical effort-based motivated task^28^. Glutamine is the main precursor for the synthesis of glutamate^35^, a building block for the production of glutathione (GSH) ^36,37^. Glutathione is a tripeptide (glutamate–cysteine–glycine) essential for several cellular functions and the most prominent antioxidant in the brain^38,39^. It seems, therefore, plausible to hypothesize that GSH levels in the NuAc may be related to the capacity to exert incentivized effort. This hypothesis is supported by convergent evidence indicating that: (i) GSH plays a major role in the reduction of reactive oxygen species (ROS) 38; (ii) ROS generation is coupled to energy production and cellular activity^40^; (iii) effortful behavior engages the activation of the NuAc^14,17–21^; and (iv) altered GSH levels have been reported in the brain of patients in disorders that course along with motivational impairments^41,42^.

To assess this hypothesis, we first explored the relationship between effort-based motivated performance in humans and accumbal levels of several metabolites, including GSH, quantified using ^1^H-MRS at 7 Tesla (7T)^28^. These experiments identified a relationship between accumbal GSH levels and motivation to exert effort. Then, to go beyond correlational evidence, we investigated the relationship between accumbal GSH levels in rats and reward-incentivized performance. To this end, we used a progressive ratio (PR) schedule of reinforcement in an operant task that requires to maintain nose-poking under increasing work demands, and modulated GSH levels through specific interventions in the NuAc. Finally, aiming at a translational application, we evaluated the efficiency of a nutritional supplementation with *N*-acetyl-cysteine (NAC) - a cysteine precursor that has been documented to increase GSH levels in the brain 39- to improve rats’ motivated performance, and explored potential effects in the excitability of the accumbal circuitry.

## Results

### Individual differences in accumbal GSH levels predict effort-based motivated performance in humans

To assess whether the concentration of GSH in the NuAc in humans is related to motivated performance, and to do so in an unbiased manner, we computed levels of a number of metabolites [creatine (Cr), glycerylphosphorylcholine (GPC), GSH, inositol (Ins), lactate (Lac), N-acetylaspartate (NAA), N-acetylaspartylglutamate (NAAG), phosphocreatine (PCr), phosphatidylethanolamine (PE) and taurine (Tau)] extracted from the ^1^H-MRS spectra acquired - but not previously analyzed- in the Strasser et al.^28^ study (Supplementary Figure 1a). Following ^1^H-MRS acquisition at 7T, participants performed an effort-based motivated task in which they could earn different monetary rewards (0.2, 0.5 or 1 CHF, depending on the trials) by squeezing a handgrip at 50% of their maximal voluntary contraction and maintaining that force for 3 sec (Figure 1a). Correlational analyses between metabolites levels and the total number of successful trials - including all rewards-indicate that, from the metabolites analyzed, GSH is the only one that shows a significant correlation (Figure 1b and Supplementary Figure 1b, c). Given that the experiment included two conditions (i.e., performance in isolation or in competition), and performance was influenced by both incentive level and competition28, we hypothesized that levels of relevant (i.e., ‘predictive’) metabolites would particularly relate to performance evoked by the higher incentive at stake (i.e., 1 CHF) for which incentive value is comparable under both social context conditions^28^. Indeed, GSH levels show a positive correlation with successful 1 CHF trials (Figure 1c), but not with performance at the lower stakes (Supplementary Figure 1b). In fact, none of the metabolites analyzed were associated with performance for lower incentive levels (Supplementary Figure 1b). Interestingly, the levels of phosphatidylethanolamine (PE), an abundant glycerophospholipid, showed as well a significant correlation with performance for 1 CHF (Figure 1b and Supplementary Figure 1d); this is an unforeseen finding that will not be followed up here, but may prompt future investigations.

**Figure 1:**
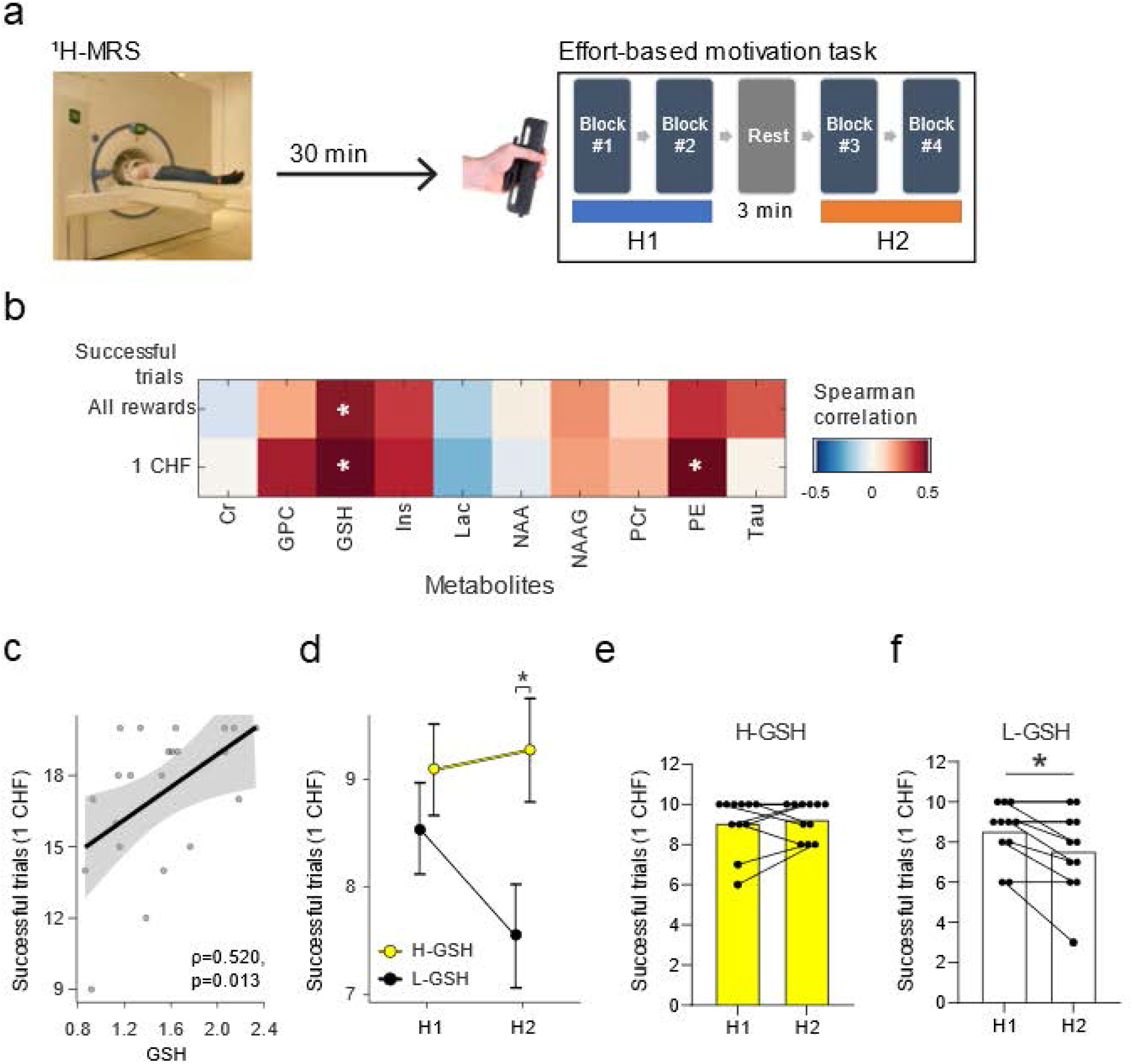
GSH levels in the nucleus accumbens are associated with the number of successful trials on an effort-based motivation task. (**a**) Experimental paradigm depiction. The ^1^H-MRS nucleus accumbens scan precedes the Effort-based motivation task, which is composed by two halves (H1 and H2), separated by a 3-minute rest block. (**b**) Matrix showing all pairwise correlations between metabolites of interest and the number of successful trials in the entire task for all rewards and for 1 CHF rewards. (**c**) Detailed scatterplot for the correlation between GSH levels in the nucleus accumbens and the number of successful 1 CHF trials (ρ=0.520, p=0.013). (**d**) Participants with lower GSH in the nucleus accumbens (L-GSH) show a lower number of successful 1 CHF trials in the second half of the experiment (H2) than those with higher nucleus accumbens GSH levels (H-GSH) (ANCOVA, H-GSH = 9.27 ± 0.91 successful 1 CHF trials wins, L-GSH = 7.55 ± 2.07 successful 1 CHF trials wins; F(1,19)= 7.52, p =0.013, η_p_^2^ = 0.284). This effect is still significant after we control for condition (isolation or competition) (F(1,18)= 7.15, p =0.015, η_p_^2^ = 0.284). Participants with high GSH levels in the nucleus accumbens are able to maintain an elevated number of successful 1 CHF trials (**e**) while those with low GSH levels in the nucleus accumbens show a significant reduction of performance in H2 (**f**) (e-f: H-GSH: t(10)=0.61, p= 0.553, Cohen’s d=0.185; L-GSH: t(10)= −3.03, p= 0.013, Cohen’s d=0.913). Significance levels: * p<0.05. Spearman correlation coefficient (ρ). N=11/group.

Then, in order to assess whether GSH levels relate to specific temporal factors in task performance, we split the data from the whole task in two parts (i.e., first and second halves: H1 and H2, each one containing 40 trials; see Figure 1a). For the analyses, we divided participants in two groups according to their accumbal GSH content (i.e., low, L-GSH: N=11; and high, H-GSH: N=11) based on a median split. As shown in Figures 1d and 1e, subjects with high accumbal GSH sustained their performance throughout the task, maintaining a similar number of successful 1 CHF trials in the first and second halves of the task. In contrast, subjects with low accumbal GSH have a significant reduction in the number of successful trials in the second half of the task compared to subjects with high accumbal GSH (Figures 1d-f). These results point to a potential role for accumbal GSH in sustaining performance over time in effortful incentivized tasks.

### Individual differences in accumbal GSH levels in rats are associated with differences in effort-based motivated performance in rats

To determine whether accumbal GSH levels may play a causal role in regulating motivated performance, we switched to rats, which are better suited for mechanistic studies. We performed experiments in males from the outbred Wistar rats as our former studies showed a key role for mitochondrial function in the NuAc in motivated behaviors in males from this strain^26,27,29^. First, using a parallel approach to our human study, we applied ^1^H-MRS at ultra-high field (9.4T) to measure rats’ GSH content in the NuAc (Figure 2a-c) and divided animals as either High or Low-GSH (i.e., H-GSH and L-GSH) following a median split, following procedures in our human study (Figure 2d). Then, we assessed rats’ willingness to exert effort in an aversively-motivated task, the forced swim test (FST), in which effort is indicated by the persistence of animal’s movement in an inescapable water-filled cylinder^43^. L-GSH rats spent more time immobile than H-GSH rats when tested in the FST over a period of 15-min (Figure 2e), with differences particularly emerging on the second half of the task (Figure 2f). The two groups did not differ in anxiety-like behaviors (Figure 2g). These results support findings from our human study establishing a link between NuAc GSH levels and motivated performance.

**Figure 2.**
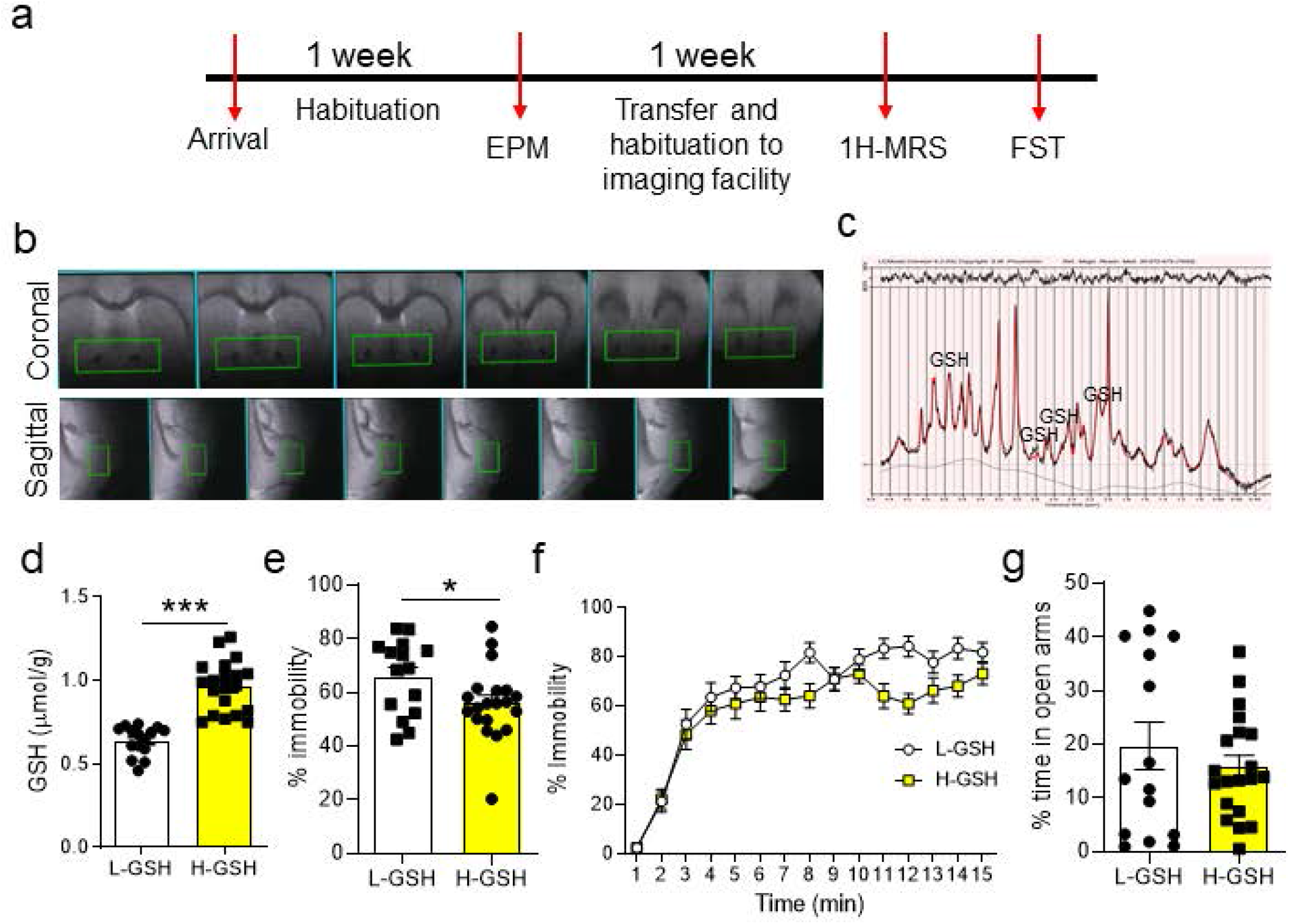
Rats differing in accumbal GSH levels show differences in immobility in the Forced Swim Test (FST). (**a**) Experimental design: Upon arrival to the facilities rats were allowed to acclimatize to the conditions and were handled to habituate to the experimenters. Then they were tested in the elevated plus maze. Subsequently, they were transferred to the imaging facilities and after a further week of habituation, they were scanned in the 1H-MRS scanner. A few days after the scanning session, rats were exposed to a 15-min FST and subsequently killed. (**b**) Representative scans from the nucleus accumbens, of animals used in the FST. (**c**) Representative GSH trace from the nucleus accumbens. (**d**) A median split was performed to split rats into two groups, High-GSH (H-GSH) and Low-GSH (L-GSH) (two-tailed t-test, t(33) = 7.272, p < 0.001). (**e**) H-GSH rats spent less time immobile in the FST (two-tailed t-test, t(33) = 2.095, p = 0.0439). (**f**) When FST performance was analyzed in 1-min bins, two-way repeated measures ANOVA revealed a significant time X GSH interaction (F(14, 504) = 2.101, p = 0.0107), a marginally non-significant effect of GSH (F(1,36) = 3.658, p = 0.0638) and a significant effect of time ((F(36, 504) = 11.87, p < 0.001), N=16-20/group. (**g**) L-GSH and H-GSH did not differ in anxiety (two-tailed t-test, t(33) = 0.8542, p = 0.3991).

In order to match our mechanistic studies in rats with the human findings reported above, we next tested rats in an effort-based reward-incentivized task, the operant conditioning-based progressive ratio (PR). This task evaluates the amount of effort an animal is willing to exert in order to obtain a sucrose pellet as a reward^44^. The level of effort at which the animal stops responding to obtain a reward is called the ‘breakpoint’^45^. Decreases in breakpoint can signal disrupted motivation. Unfortunately, our neuroimaging facility is not equipped with operant boxes and its sanitary status implies that rats cannot be moved after ^1^H-MRS testing back to the main animal facility where the behavioral tasks are located. Thus, given this technical constrain, in the next experiments accumbal GSH content was quantified *post-mortem.*

Thus, a new cohort of rats was trained in an operant conditioning paradigm, involving a 6-day fixed ratio 1 (FR1) training followed by the PR test (Figure 3a and 3b). GSH was measured using HPLC following the harvest of the NuAc (Figure 3C), which allowed establishing H-GSH and L-GSH groups (Figure 3d). The two groups showed no differences in anxiety-like behaviors in the elevated plus maze (EPM) or time spent in the center of Open Field (OF) (Supplementary Figure 2 a, b), in agreement with our results obtained using 1H-MRS (Figure 2g). The groups also did not differ in locomotion (Supplementary Figure 3c) or exploration (Supplementary Figure 2c) as measured in OF and novel object (NO) tests, respectively. Moreover, the two groups did not differ in their performance with a low-effort fixed ratio schedule (Supplementary figure 2e), indicating that differences in accumbal GSH levels are not related to differences in learning or in the ability to work for rewards in a low-effort task. However, in the increasing effort-based PR task, H-GSH rats exhibited a higher breakpoint (Figure 3e), obtained more rewards (Figure 2f) and performed more correct nose pokes (Figure 3g) than L-GSH rats. Importantly, the two groups did not differ in the percentage of correct over total (correct and incorrect) nose pokes (Supplementary Figure 2f). This indicates that the observed group differences in correct nose pokes are not due to differences in accuracy of responses or in randomly responding to both the correct and incorrect ports, which could have indicated differences in goal-directed behavior. Survival curve analyses showed that a significantly higher number of H-GSH rats reached higher breakpoints (Supplementary Figure 2g) and this group also showed a trend towards nose poking over a longer period of time (Supplementary Figure 2h). No group differences were observed when analyzing rats’ responses over the different ratio requirements (Supplementary Figure 3i), but analysis of the number of correct nose pokes over time revealed a significant effect of time and GSH levels (Supplementary Figure 2j). As in humans, our animal model indicates a role of accumbal GSH in the ability to perform as well as to persevere in an effortful incentivized task. Finally, when subsets of H-GSH and L-GSH rats were allowed free access to 20 sucrose pellets (more than any rat managed to acquire during the time they are given access to the PR test), all animals readily consumed all pellets (Supplementary Figure 2k), suggesting no group differences in appetite or in the perceived palatability of the pellets.

**Figure 3.**
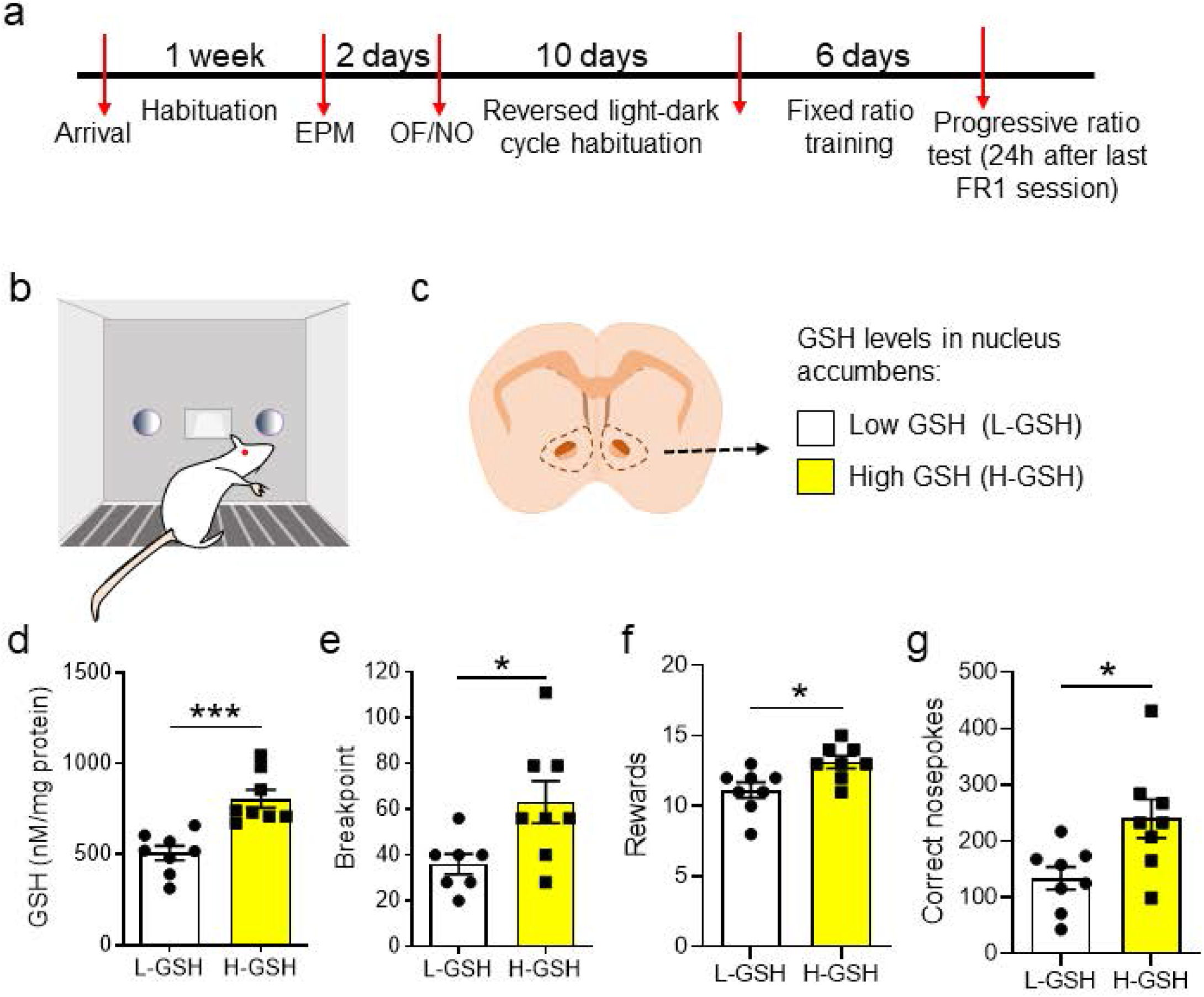
High GSH levels in nucleus accumbens are associated with an improvement in Progressive Ratio (PR) test performance. **(a)** Timeline showing the experimental design. After initial acclimatization, handling and behavioral characterization for anxiety and exploratory behavior, rats were placed in reversed light-dark cycle to habituate for ten days. Subsequently, rats were trained in the Fixed ratio 1 paradigm for a total of six sessions (depicted in **b**). Once the PR test was performed, rats were sacrificed, their nucleus accumbens was microdissected and snap frozen on liquid nitrogen. Accumbal GSH levels were measured using HPLC (scheme depicted in **c**). (**d**) Rats were split into two groups based on their accumbal GSH levels (High GSH and Low GSH; H-GSH and L-GSH, respectively), using a median split (t(14) = 4.686, p < 0.001) In the PR test, H-GSH rats reached a higher breakpoint (t(14) = 4.686, p < 0.001) (**e**), obtained more rewards (t(14) = 2.841, p = 0.0131)(**f**) and performed more correct nosepokes (t(14) = 2.660, p = 0.0187) (**g**) indicating that animals with high levels of GSH in nucleus accumbens performed better in the PR paradigm. Unpaired t test two-tailed (***p<0.0001, * p<0.05, N=7-9/group).

Taken together, these data indicate that H-GSH rats have a longer active engagement in the PR test than L-GSH rats and, their superior performance is not due to a higher rate of responses at the beginning of the PR test. These results indicate a higher endurance capacity to work for rewards over time for rats with higher accumbal GSH levels, resembling the data reported above for human subjects. Therefore, we selected this test and approach to investigate the causal involvement of accumbal GSH in motivated performance.

### Administration of a GSH inhibitor into nucleus accumbens impairs PR test performance

To assess whether decreasing local accumbal GSH levels alters motivated performance, rats were injected with either BSO (an inhibitor of gamma-glutamylcysteine synthetase-the rate-limiting enzyme for GSH synthesis) at 7 μg/μl or DMSO (the diluent of BSO) intra-NuAc (1 μl per hemisphere) (Figure 4a and 4b). Both groups were matched for anxiety, exploratory and locomotor behaviors (before cannulation) and performance during the fixed ratio training (Supplementary Figures 3a-c). Following training, each group of rats was microinfused with BSO or DMSO 24 hours prior to the PR test. Our BSO dose was effective in reducing GSH levels as verified via HPLC measurements (Figure 4c). In the PR test, BSO-infused animals achieved a lower breakpoint (Figure 4d), obtained fewer rewards (Figure 4e) and performed fewer correct nosepokes (Figure 4f) than the DMSO group. No group differences were observed in the percentage of correct nosepokes over the total number of nosepokes (Supplementary Figure 3d). BSO-treated rats had a lower probability to reach a high breakpoint (Supplementary Figure 3e) and a non-significant tendency to perform fewer nosepokes (Supplementary Figure 3f) than the DMSO group. Performance over the different ratio requirements was also not significantly different between the two groups (Supplementary Figure 3g). However, BSO-treated rats showed a significantly lower number of correct nosepokes over time (Supplementary Figure 3h).

**Figure 4.**
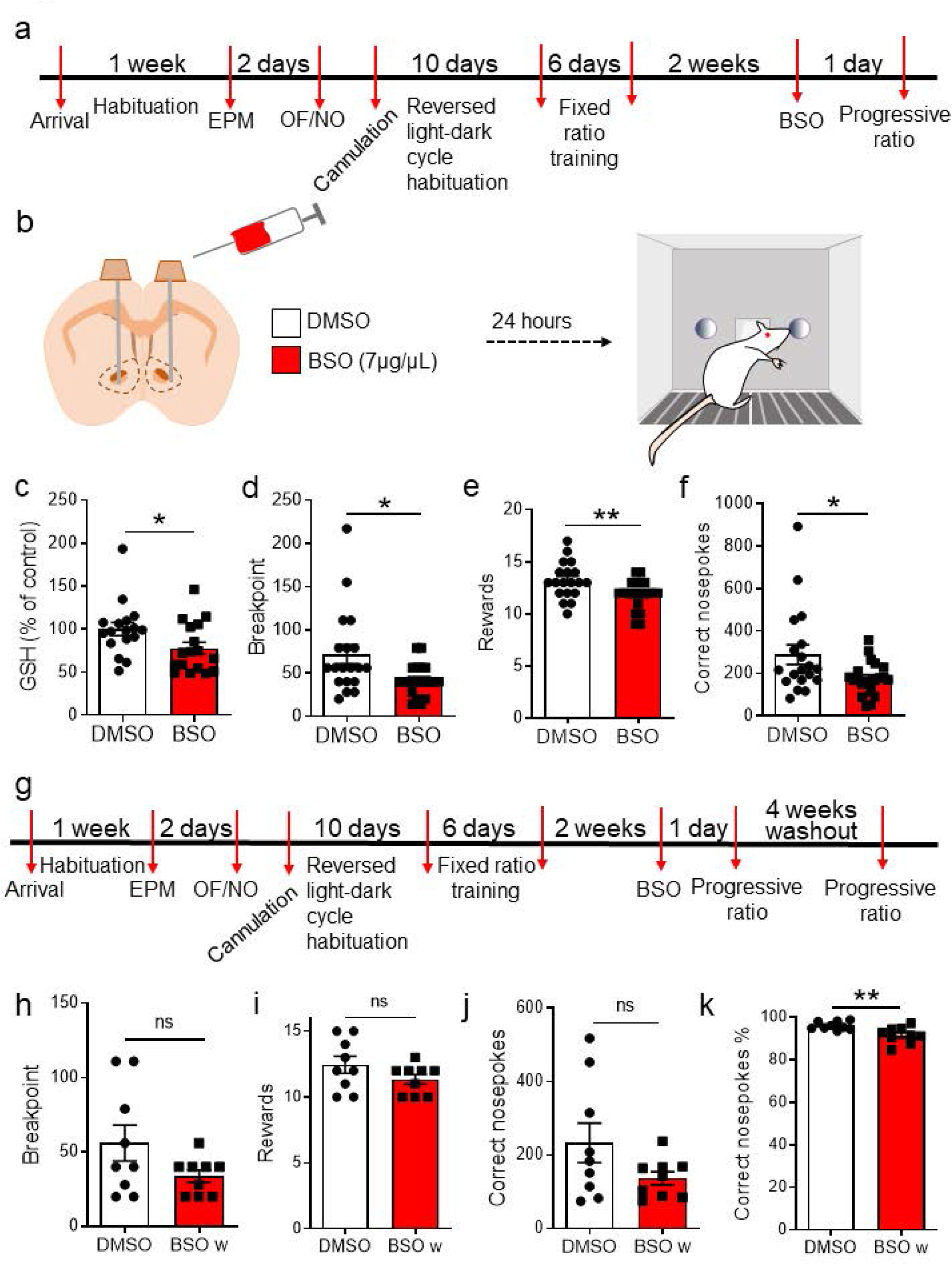
Intra-accumbal administration of BSO impairs PR performance. (**a**) Cannulated rats habituated for 10 days in reversed light-dark cycle during their recovery from surgery. Subsequently, they were trained in an FR1 schedule for a total of six sessions. (**b**) 24 hours before the PR test, BSO or DMSO was infused via a cannula into the nucleus accumbens. (**c**) GSH levels were measured in the nucleus accumbens of DMSO- and BSO-infused rats by HPLC demonstrating the ability of BSO to reduce GSH levels (unpaired two-tailed t-test t(32) = 2.154, p = 0.0389 N=17/group). (**d**) PR performance was impaired in BSO infused animals compared to the vehicle-treated group, as indicated by the lower breakpoint (unpaired two-tailed t-test t(38) = 2.644, p = 0.0119). (**e**) The number of obtained rewards was also lower following intra-accumbal BSO treatment (unpaired two-tailed t-test t(38) = 2.722, p = 0.0097). (**f**) BSO-treated rats performed fewer correct nosepokes compared to vehicle-treated rats (unpaired twotailed t-test t(38) = 2.376, p = 0.0226, N = 19-21/ group). (**g**) Timeline showing the experimental design for the washout experiment. Rats previously treated with vehicle or DMSO were tested again in the PR task, four weeks after the previous test. (**h**) The breakpoint was not different between groups (unpaired twotailed t-test, t(16) = 1.751, p = 0.0992). (**i**) The number of obtained rewards was not significantly different between groups (unpaired two-tailed t-test, t(16) = 1.487, p = 0.156). (**j**) The number of correct nosepokes was not significantly different between groups (unpaired two-tailed t-test, t(16) = 1.704, p = 0.108). (**k**) Rats previously treated with BSO performed a lower percentage of correct nosepokes, over the total number of nosepokes in the washout session (unpaired two-tailed t-test, t(16) = 3.133, p = 0.0064, N = 9 per group).

To rule out a possible toxic effect on the NuAc of the BSO dose used, we performed an activated caspase 3 (a marker of the execution-phase of cellular apoptosis) and neuroN (a marker of mature neurons) staining assay on accumbal sections from a subset of animals tested in the PR test (Supplementary Figure 3 i-l). No staining differences were observed between DMSO and BSO groups, indicating a lack of BSO toxicity at the dose used in this experiment.

To further confirm that the effects of BSO on motivated performance are not due to any sort of toxicity or profound alteration of circuit function, after a first PR test, a subset of BSO- and DMSO-infused animals underwent a 4-week washout period and were then submitted again to the PR test (Figure 4g). Importantly, 4 weeks after BSO treatment, no differences between BSO- and DMSO-treated rats were observed on the breakpoint reached (Figure 4h), the number of obtained rewards (Figure 4I) or the number of correct nosepokes (Figure 4j). BSO-washout animals exhibited a small, but significant, decrease in the percentage of correct nosepokes over all nosepokes (both in the active and the inactive port) (Figure 4k) despite the lack of significant differences in correct nosepokes performed after the washout (Figure 4j). These set of data supports the view that lowering accumbal GSH levels leads to a transient impairment in effort-based incentivized performance without inducing toxicity.

### N-acetyl-cysteine (NAC) treatment increases accumbal GSH levels and improves PR test performance

Next, we investigated the converse approach; i.e., enhancing GSH synthesis to assess the impact on PR performance. To this end, rats were trained in a FR1 schedule and then segregated in two equivalent groups matched according to their performance at training (Supplementary Figure 4a), body weight (Supplementary Figure 4b) and anxiety levels (Supplementary Figure 4c). Then, one of the groups received NAC in the drinking water at a concentration of 0.5 mg/L for two weeks while the other group received normal drinking water (Vehicle) (Figure 5a and b). NAC treatment provides a source of cysteine, a rate-limiting building block for GSH synthesis, and is known to increase GSH levels in the brain^46^. Here we confirmed its effectiveness to increase accumbal GSH levels in a separate cohort of animals (Figure 5c) in which we also confirmed no drastic changes in basal levels of anxiety-like behaviors (Supplementary Figures 4d, e). NAC-treated rats performed better in the PR session than vehicle-treated rats, achieving a higher breakpoint (Figure 5d), obtaining more rewards (Figure 5e) and performing more correct nosepokes (Figure 5f). The percentage of correct nosepokes over the total number of nosepokes (both correct and incorrect) was not different between the two groups (Supplementary Figure 4f). Survival analysis showed that a higher percentage of NAC-treated rats reached higher breakpoints (Supplementary Figure 4g) and kept nose poking for longer (Supplementary Figure 4h). NAC treatment did not affect rats’ performance over the different ratio requirements (Supplementary Figure 4i) or the number of correct nosepokes over time (Supplementary Figure 4j). Altogether, these data show that a 2-week supplementation with NAC in the drinking bottle increases GSH levels in the NuAc and improves rewardbased effortful performance and endurance.

**Figure 5.**
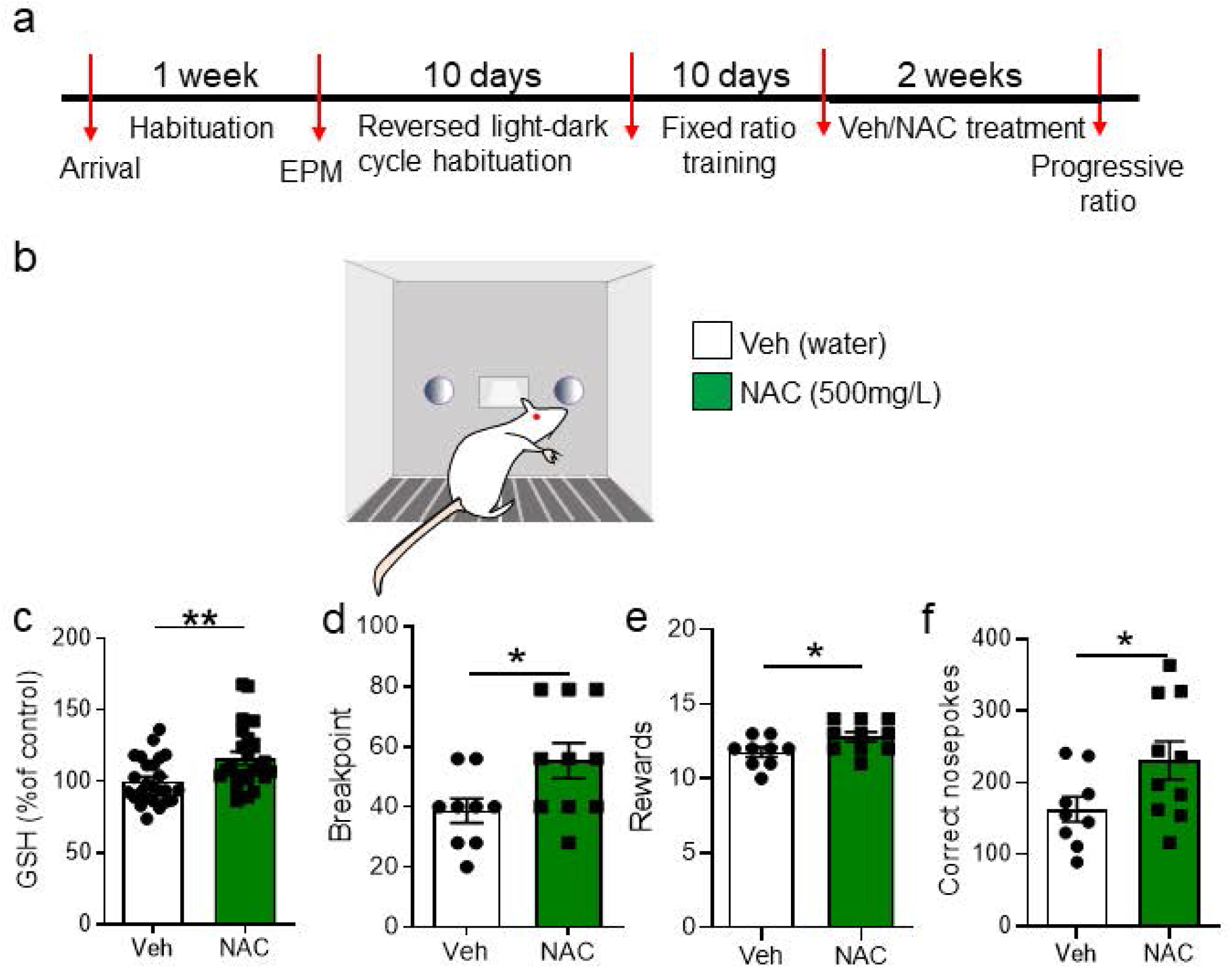
NAC systemic treatment increases GSH levels in nucleus accumbens and ameliorates PR task accomplishment. (**a**) Timeline of the experiment. After initial habituation, handling and behavioral characterization for anxiety, rats were placed in reverse light-dark cycle and trained in an FR1 schedule. Once matched for performance and weight, rats were distributed into Vehicle and NAC groups and received normal drinking water or 500 mg/L of NAC in the drinking water, for two weeks. Then, both groups were tested in the PR test (**b**). (**c**) A separate cohort of Vehicle- or NAC-treated animals was sacrificed and GSH levels were measured in the nucleus accumbens (unpaired two-tailed t-test, t(47) =2.968, p = 0.0047, N=24-25/group). (**d**) NAC-treated rats reached a higher breakpoint than vehicle-treated rats (unpaired two-tailed t-test, t(17) = 2.271, p = 0.0364). (**e**) Similarly, NAC-treated rats obtained more rewards than vehicle-treated rats (unpaired two-tailed t-test, t(17) = 2.215, p = 0.0407). (**f**) NAC-treated rats performed more correct nosepokes than vehicle treated rats (unpaired two-tailed t-test, t(15.24) = 2.094, p = 0.0516) (**p<0.01, * p≤0.05, N=9-10/group).

### NAC treatment affects excitatory inputs onto MSNs in a cell-type specific manner

Next, we asked whether the improvement in behavioral performance induced by NAC treatment was accompanied by changes in the excitability of the accumbal circuitry. The core division of the NAc is essential for cue-motivated behaviors such as performance in the PR task^47^. Its two main neuronal populations [i.e. medium spiny neurons expressing the dopamine receptor type 1 (D1-MSNs) and type 2 (D2-MSNs)] are classically described as mediators of distinct types of behavior (e.g., approach versus avoidance)^48,49^. We performed *ex vivo* patch-clamp recordings in the NAc core from NAC- and vehicle-treated rats and identified the cell type of the recorded cells by biocytin-filling and *post hoc* in-situ hybridization (Figure 6a-c). No differences were observed in passive cell properties, spiking threshold and firing frequency when cells were depolarized by somatic current injections (Supplementary Figure 5a-d), indicating that NAC treatment did not affect intrinsic excitability of MSNs. However, we observed celltype specific alterations in the excitatory inputs. Miniature postsynaptic excitatory currents (mEPSCs) recorded in D1-MSNs exhibited significantly larger peak amplitudes in NAC-treated rats, with no differences in the inter event interval (Figure 6d). By contrast, mEPSC peak amplitudes were significantly reduced in D2-MSNs, and were also less frequent, as indicated by the pronounced increase in the interevent interval (Figure 6e). Overall, these data indicate that the weight of glutamatergic synaptic drive in the NAc core is altered by the NAC treatment, such that D1-MSNs receive larger excitatory inputs, while these are overall reduced in D2-MSNs. The resulting augmented engagement of D1-MSN over D2-MSN is consistent with the purported roles of the two cell types in approach/avoidance behavior, thus offering a plausible cellular substrate for the enhanced performance in the PR task.

**Figure 6.**
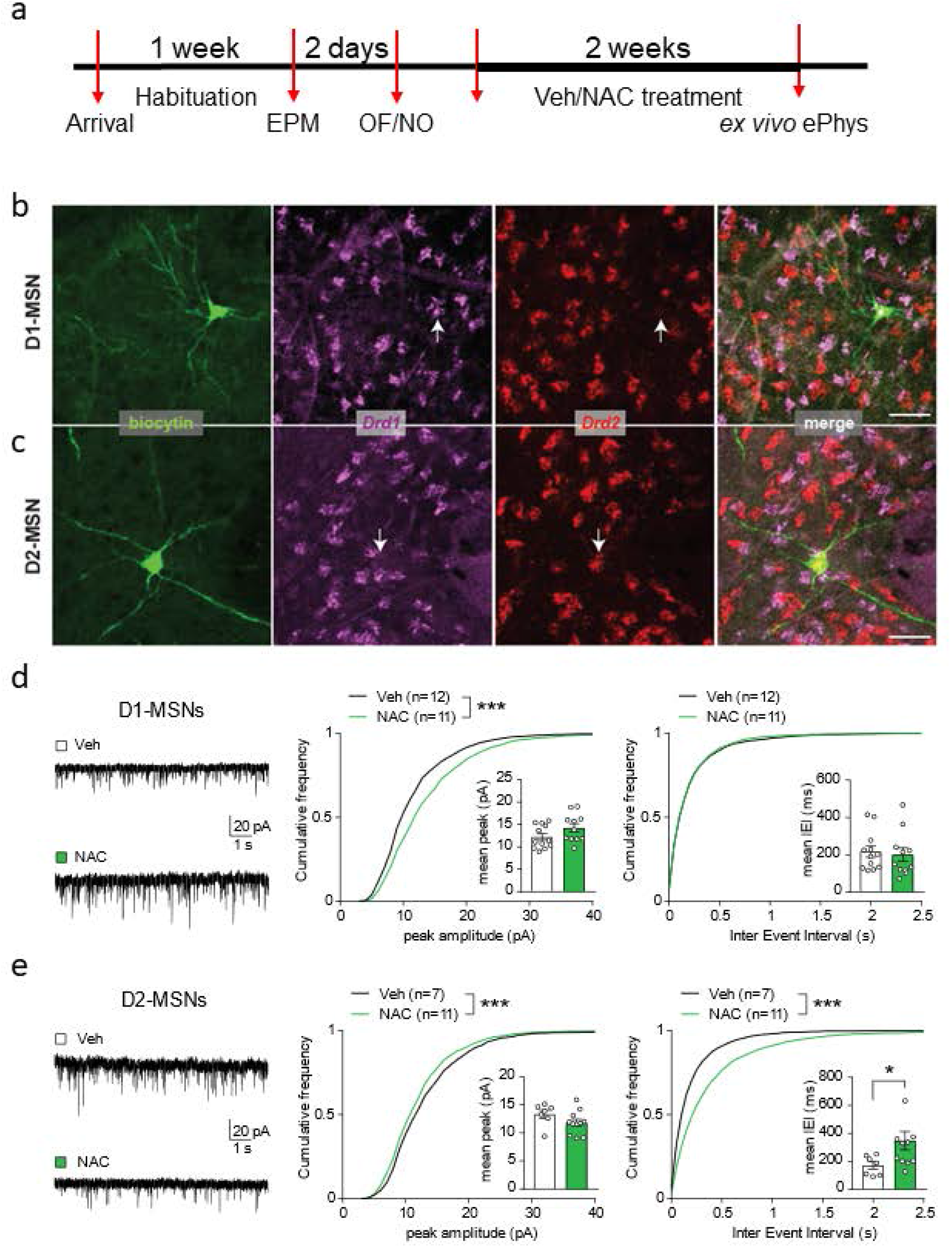
NAC treatment affects excitatory inputs onto MSNs in a cell-type specific manner. (**a**) Experimental timeline for ex vivo electrophysiological recordings in NAC-treated rats. (**b-c**) Confocal micrographs showing neurons in the nucleus accumbens core filled with biocytin during patch-clamp recordings and identified *post hoc* as D1-MSN (**b**) and D2-MSN (**c**) based on RNAscope reactivity for *Drd1* and *Drd2* mRNA (color-coded). Scale bar, 50 μm. (**d**) Left, example traces of miniature excitatory postsynaptic currents (mEPSCs) in D1-MSNs from vehicle- and NAC-treated rats. Cumulative frequency plots indicate larger peak amplitude in NAC-treated rats (middle panel, Kolmogorov-Smirnov test, D=0.1611, ***p<0.001), with no change in mEPSC inter event interval (right panel, Kolmogorov-Smirnov test, D=0.0181, p=0.8453). Insets with bar graphs represent mean values per cell (peak, unpaired t test, t=1.691, p=0.1067; inter event interval (IEI), Mann-Whitney test, U=56, p=0.5658). (**e**) Left, example traces of mEPSCs in D2-MSNs from vehicle- and NAC-treated rats. Cumulative frequency plots indicate smaller peak amplitude (middle panel, Kolmogorov-Smirnov test, D=0.0752, ***p<0.001) and larger inter event interval in NAC-treated rats (right panel, Kolmogorov-Smirnov test, D=0.2213, ***p<0.001). Insets with bar graphs represent mean values per cell (peak, unpaired t test, t=1.586, p=0.1354; IEI, Mann-Whitney test, U=16, *p=0.0441).

## Discussion

Here, we establish a novel role for GSH in specific brain structures in motivation. Specifically, we identify GSH levels in the NuAc as critically related to the individual’s capacity to keep on exerting effortful behavior over time in reward-incentivized tasks. We first underscored GSH levels in the NuAc as predictive of steady execution in an effort-based monetary incentivized task in humans. Participants with higher accumbal GSH levels were able to steadily maintain good performance until the end of the task, while those with lower levels showed a decay in the second half. To investigate causality, we switched to rats in which we confirmed that individual differences in accumbal GSH levels predict performance in both aversively- and appetitively-motivated tasks. The involvement of GSH in effort-based motivated performance was, supported by the observation of impaired performance in the PR test following down-regulation of accumbal GSH levels through the microinjection of the GSH synthesis inhibitor BSO. Furthermore, systemic treatment with NAC, a source of cysteine known to increase brain GSH, led to increases in accumbal GSH levels as well as in rats’ breakpoint in the PR task, that was possibly due to a shift in glutamatergic inputs to D1- and D2-MSNs. Our results highlight the ability of accumbal GSH levels to facilitate motivated endurance over time. These findings may open new avenues to consider accumbal GSH as both a biomarker for motivational disturbances and a potential target for therapeutic interventions.

Although we set the study on the specific hypothesis that accumbal GSH may be involved in effort-based motivated performance, our first unbiased analysis considering a range of metabolites measured through ^1^H-MRS at 7T in humans yielded GSH as the metabolite specifically predictive of performance out of a total of 10 metabolites analyzed. Given that GSH shows a low physiological concentration and its spectra strongly overlaps with other metabolites, its measurement is challenging especially at low magnetic fields. Increasing spectral signal-to-noise ratio and resolution at high magnetic field (7T and above) as used here is critical for improving metabolites quantification especially for J-coupled metabolites such as GSH ^50,51^. Using a similar approach in rats (^1^H-MRS at 9.4T), we confirmed the capacity of accumbal GSH levels to predict performance in an aversively-motivated task, and *post-mortem* analyses related GSH levels to performance in a reward-incentivized task. Specifically, rats with higher accumbal GSH levels spent less time immobile in the forced swim test (i.e., displaying more active coping responses to adversity) and had higher breakpoints in the PR task than their lower GSH counterparts.

Formerly, we identified a predictive value for glutamine, and particularly the glutamine-to-glutamate ratio in motivated performance in humans, and hypothesized the contribution of these metabolites to fuel mitochondrial function under enhanced NuAc engagement during effortful incentivized behaviors. Indeed, human studies have shown that the NuAc (or the ventral striatum more broadly) is the main brain structure to respond to incentives^14,16,52,53^ being incrementally recruited as the expected reward increases ^16,54^. Nucleus accumbens activity relates to effort-based cost-benefit valuation^12^, acting as the motivating force^17^ that engages both cognitive and motor downstream brain regions^14^. These human studies measured blood oxygen level dependent (BOLD) signals through functional magnetic resonance imaging (fMRI), as a proxy of neural activity. Work in rats using *in vivo* oxygen (O_2_) amperometry similarly showed increased O_2_ responses to rewards as a function of increased required effort to obtain them. Animals with higher breakpoints showed a greater magnitude of the NuAc O_2_ responses, suggesting that individual differences in the capacity to remain on task for longer are related to the magnitude of O_2_ responses in the NuAc^18^. Given that the main O_2_-consuming cells in the brain are neurons^55^, increased O_2_ in the NuAc mainly represents the energetic cost of neuronal activity. Specifically, oxygen consumption enables to meet increased energy demands in neural circuits by facilitating oxidative phosphorylation in the mitochondria and thus, contributing to the generation of adenosine triphosphate (ATP)^14,55,56^. However, oxygen consumption generates free radicals and other reactive oxygen species which, when excessive and not sufficiently counteracted by cellular antioxidant systems, can produce oxidative stress, toxic to cells and molecules, and to which the brain is particularly susceptible^40^. Importantly, higher mitochondrial oxygen respiratory capacity in the NuAc in rodents has been shown to positively related to the duration of active coping/escaping responses in the FST^29^ and to winning a social competition^26,27^. Accordingly, our findings here suggest that a higher accumbal GSH production may act in concert with mitochondrial oxidative phosphorylation processes to facilitate motivated effort. Its causal contribution to motivated performance was established in our study by the direct inhibition of GSH levels by BSO intra-NuAc injection.

Glutathione is the major antioxidant as well as a redox and cell-signaling regulator^38,57^. Given the high susceptibility of the brain to oxidative stress^40^, neurons require high levels of GSH to counteract oxidative stress and sustain cellular integrity and circuit function^58^. Therefore, our results, in both humans and rats, relating individual differences in accumbal GSH content to the capacity to succeed in task performance over time, supports the view that higher individual GSH levels provide an advantage to deal with oxidative processes generated by the drastically enhanced energy demand - involving increased O_2_ utilization-required in NuAc circuits to maintain performance (see above). However, given our previous findings implicating the glutamine-glutamate ratio in effortful performance^28^, and recent evidence indicating that GSH may be a physiological reservoir of glutamate neurotransmitter when the glutamate-glutamine shuttle is inhibited^59^, the alternative non-mutually exclusive possibility that higher GSH levels may be contributing to the glutamate-glutamine cycle [or *vice versa*; note that glutamine may also contribute directly to GSH synthesis^37^] and, thus, facilitating performance cannot be ruled out.

We also carried out a translational approach and examined the potential of a prolonged treatment with the GSH booster NAC to improve motivation. Our administration regime effectively increased accumbal GSH content, supporting the efficiency of systemic NAC treatment in targeting antioxidant levels in the ventral striatum. Importantly, NAC treatment was also effective in increasing rats’ breakpoint in the PR task, supporting its ability to ameliorate motivational deficiencies. Selective activation of specific glutamatergic inputs to the NuAc has been shown to promote reward-based motivation^60–62^. Importantly, we provide novel evidence that prolonged systemic treatment with NAC induces cell subtype-specific changes in synaptic transmission in the NuAc core: quantal glutamatergic currents were larger in D1-MSNs, while the excitatory drive onto D2-MSNs was overall diminished. This suggests that, in NAC-treated rats, excitatory inputs promote activation of D1-MSNs over D2-MSNs as a result of the increased postsynaptic responses in D1-MSNs as well as the concomitant reduction in collateral inhibition provided by D2-MSNs^63^. The shift towards D1-MSN activation is expected to support motivated behavior, according to the canonical roles of D1- and D2-MSNs in positive reinforcement and aversion, respectively [^47–49^; but note that several studies have suggested that D2-MSNs may also contribute to reward-based motivation^63–65^ and that MSN subtype-specific actions may depend on different patterns of activation^66^]. The mechanisms whereby NAC treatment can affect glutamatergic function in the NuAc are multiple. In addition to promoting GSH synthesis, cystine derived from NAC also binds the cystine-glutamate exchanger in glial cells and shifts the extracellular levels of glutamate^67^. Moreover, alterations in the metabolic pathways leading to the synthesis of GSH can affect glutamate levels and accumulation^68^ and glutamate activity can also be shaped by GSH^59^. Future studies are needed to tackle the molecular mechanisms mediating these cell-type specific rearrangements. Notably, opposite effects on glutamatergic transmission have been shown to occur in D1- and D2-MSNs upon drug exposure^69^, confirming that synaptic adaptations in the NuAc can take place in a cell-type specific manner. NAC interventions have been applied in animal models of addiction and have proven successful in rescuing drug-seeking behavior by normalizing extracellular glutamate levels, thereby modulating accumbal excitatory transmission via metabotropic receptors^70^. However, in contrast to our results, such cellular mechanisms have been addressed in response to acute NAC administration and do not exhibit cell-type specificity. Thus, our findings are likely the result of the potent antioxidant capacity of GSH facilitating the engagement of accumbal circuitry to produce motivated behavior, as well as GSH-independent mechanisms involving more complex and long-term modifications, including a reshaping of synaptic glutamatergic activity.

Variability in accumbal GSH levels can be due to both genetic and non-genetic life and situational factors. Importantly, as shown in some cell types, intra-individual variation in GSH content seems to be rather stable over time^71^. Individual capacity to produce GSH is determined by genetic variability in enzymes involved in its production and/or regeneration, including several common single nucleotide polymorphisms (SNPs) that regulate GSH levels and associated processes^72^. In addition, several life conditions can also affect brain GSH levels, particularly under situations leading to increased ROS expression and oxidative stress such during ageing or psychogenic stress^39,73^. Accordingly, several psychopathological alterations - such as depression or schizophrenia-, in which low GSH levels have been reported, are associated with motivational deficits or fatigue^41,42,74^. Remarkably, we report that interindividual differences in accumbal GSH levels in healthy, human and rodent populations (i.e., within the normal physiological range) are related to differences in effortful behavior. Therefore, our findings here go well beyond previously reported correlations of brain GSH levels with psychiatric conditions to causally implicate accumbal GSH levels in motivated behavior in healthy individuals. Moreover, we provide a proof of principle that nutritional treatments leading to 15-20% change in accumbal GSH content are able to modulate motivated performance in healthy individuals.

In conclusion, we establish GSH levels in the NuAc as both a predictive marker of differences in reward-based effortful performance and as a potential target for nutritional or other type of interventions. We also provide strong evidence for a promising potential of chronic NAC supplementation to boost accumbal GSH levels and to regulate motivation to exert reward-incentivized effort.

## Materials and Methods

### Human study: Participants

Forty-three men, between 20–30 years old, were originally recruited for the study. Informed consent was obtained from all participants, who were debriefed at the end, and experiments were performed in accordance with the Declaration of Helsinki and approved by the Cantonal Ethics Committee of Vaud, Switzerland. Valid NuAc ^1^H-MRS and behavioral data were obtained from 22 participants (more experimental details can be found in the Participants section in Supplementary Methods of ref^28^). Data analyzed here have not been previously analyzed but it were collected as part of a larger study previously published^6,28^. Ethics: The experiment was performed in accordance with the Declaration of Helsinki and approved by the Cantonal Ethics Committee of Vaud, Switzerland.

### Effort-related monetary incentivized force task

Our task, termed monetary incentive force (MIF) task^6,28^, combines aspects from the monetary incentive delay (MID) task^16^ and from effort-based decision-making paradigms^10,17^. It requires participants to exert force on a hand grip dynamometer (TSD121B-MRI, Biopac) (Figure 1a) at a threshold corresponding to 50% of each participant’s maximum voluntary contraction (MVC). The experiment was run under two experimental conditions, with half of the participants performing the task in isolation (i.e., their earnings depended solely on their own performance) while the other half in competition. Success was measured by the number of successful experimental trials, and results were computed separately for the two halves of the experiment (H1 and H2; see Figure 1a) for three monetary incentives (i.e., 0.2, 0.5, and 1 CHF). See^28^ for further details.

### Proton magnetic resonance spectroscopy (^1^H MRS) acquisition for participants and data processing

The MR measurements were performed on a Magnetom 7T/68-cm head scanner (Siemens, Erlangen, Germany) equipped with a single-channel quadrature transmit and a 32-channel receive coil (Nova Medical Inc., MA, USA). The NuAc region of interest voxel (VOI) was defined by the third ventricle medially, the subcallosal area inferiorly, and the body of the caudate nucleus and the putamen laterally and superiorly, in line with definitions of NuAc anatomy identifiable on MRIs [34] See^28^. The Cramer-Rao Lower Bound (CRLB) for GSH was 8.2 +/− 2.8%.

### Animals

Adult male Wistar rats (Charles Rivers, Saint-German-Nuelle, France) weighing 250-275 g upon arrival were individually housed with *ad libitum* access to food and water in a 12 light/dark cycle (lights switched on at 7:00 am) and a constant temperature at 22 ± 2°C. Following a week of acclimatization to the animal facilities, rats were handled for 2 min/ day for three days prior to the start of the experiments, in order to habituate to the experimenters. Rats used for operant conditioning experiments were placed in reversed light-dark cycle (lights-on at 20:00, lights-off at 8:00) with *ad libitum* food and water and such experiments were performed during the dark phase of the cycle. Experiments were carried out in accordance with the European Union Directive of 22^nd^ September 2010 ([Directive 2010/63/EU](https://eur-lex.europa.eu/eli/dir/2010/63/2019-06-26)) and approved by the Cantonal Veterinary Authorities (Vaud, Switzerland). All efforts were done to minimize animal number and suffering.

### Anxiety classification

All rats were tested in the elevated plus maze (EPM) to phenotype for anxiety, as previously described^26^. Briefly, the EPM consisted of two open and two closed arms (45 x 10 cm each) extending from a 10 x 10 cm central area. Closed arms had 10 cm high walls. Lighting was maintained at 16-17 lx in the open arms, 10-11 lx in the central area and 5-7 lx in the closed arms. Rats were placed on the central area, facing a closed arm and were allowed to explore freely for 5 min. Every animal was recorded and tracking was performed using Ethovision software (Noldus, Wageningen, the Netherlands). Percentage of time in the open arm was calculated using Ethovision (Noldus).

### Open Field (OF)

The open field was used to determine locomotor behavior and time spend in the center as a measure of rat’s exploration. The open field apparatus and procedure were previously described^75^. Briefly, the open field consisted of a black circular arena (1 m in diameter, surrounded by walls 32 cm high). For analysis, the percentage of time in the centre and total distance walked were calculated. Animals were placed in the center of the arena and their behavior was monitored for 10 min using a video camera that was mounted from the ceiling above the center of the arena. The light was adjusted to a level of 8-10 lx in the center of the arena. Total distance walked and % time in the center were calculated using Ethovision (Noldus).

### Novel object (NO)

Immediately after the open field test, rats were exposed to a novel object (NO). For this purpose, a small, plastic bottle was placed into the center of the open field while the rat was inside. Rats were then given 5 min to freely explore the novel object. The time spent exploring the novel object was recorded using Ethovision and plotted as a percentage of time with the object.

### Proton magnetic resonance spectroscopy (^1^H-MRS) for rats

After anxiety and locomotion characterization, rats were transferred to the Center for Bioimaging (CIBM), EPFL, where the *in vivo* proton magnetic resonance spectroscopy (^1^H-MRS) measurements of brain metabolites was performed. Briefly, all experiments were performed on a horizontal 9.4 T/31 cm Bore animal MR scanner (Magnex Scientific, Abingdon, UK) using a homemade quadrature ^1^H-coil. Animals were anesthetized and placed in the scanner. Acquisition was done using the spin echo full intensity acquired localized (SPECIAL) sequence in the volume of interest (VOI) placed in the bilateral NuAc (voxel size: 2 × 6 × 3 mm^3^, TE/TR: 2.8/4000 ms) after acquisition of a set of anatomical T2-weighted images for localization. Field homogeneity was adjusted using FAST(EST)MAP to reach a typical water linewidth of 13.2 Hz in NuAc. Spectra were acquired with 10 blocks of 25 averages, leading to a scan duration of around 30 min respectively. After post-processing of the spectra, metabolite concentrations as well as the Cramér-Rao lower bounds (CRLB) were determined with LCModel using water as internal reference^31^.

### Forced Swim test (FST)

The FST was performed for 15 min under light conditions of 60 lx. Rats were individually placed into glass cylinders (20 cm diameter, 46cm depth) containing water at 23-25°C. The cylinder was half filled, in order to prevent the rat from touching the bottom of the cylinder with its feet or tail. The water was changed after each animal’s session and the cylinder cleaned. The percentage of Immobility time was scored manually using the Observer software (Noldus) by an experimenter blind to each animal’s allocation to a group.

### NAC treatment

For experiments including NAC treatment, rats were treated with 500 mg/L in the drinking water, for at least two weeks before the beginning of behavioral experiments. Water bottles were changed twice per week to ensure NAC integrity. For operant experiments, treatment was provided starting on the last fixed ratio training day and continued until the end of experiments.

### Stereotactic surgery for cannula implantation

Rats were anesthetized by isoflurane inhalation (4 %, for 4 min) in an induction chamber and maintained afterwards with 2 % isoflurane with a flow of 4 l/min. Stereotactic surgery was performed as previously described^26,76,77^. Briefly, rats were mounted on a stereotactic frame (Kopf Instruments, Tujunga, CA, US), an incision was made along the midline of the skull, the periosteum was removed and small holes were drilled for the implantation of guide cannulae (Invivo1, Roanoke, VA, USA). Coordinates for NuAc were taken from the Paxinos and Watson brain atlas, relative to bregma, as follows: Anterior-posterior: +1.2, mediolateral: ±1.5, dorsoventral: −6.50. Cannulae were fixated on the skull with three anchoring screws and Paladur acrylic dental cement (Kulzer, Hanau, Germany). Correct cannula placement was confirmed in the end of experiments.

### BSO infusions

Behavioral experiments were performed 24 hours after BSO or saline (Sal) administration. Groups were randomized. BSO was dissolved in DMSO at a concentration of 7 μg/μl. For local intra-cerebral infusions, the dummy cannulae were removed and injectors were inserted extending 2 mm from the guide cannulae. Drugs were bilaterally infused intra-VTA at a volume of 1 μl per hemisphere at a rate of 0.3 μl/min. The injector remained in place for one additional minute after infusion to allow proper diffusion.

### Progressive ratio task

After at least 10 days after surgery or after introduction to the reversed light-dark cycle, animals started training in a fixed ratio 1 reinforcement schedule (FR1). Operant chambers (Coulbourn Instruments, Holliston, MA, US), placed in sound attenuating cubicles, were equipped with a grid, underneath which a tray with standard bedding material was placed for collection of faeces and urine after each training session. Each chamber had one food tray and two ports placed on either side of the tray. A cue light was placed in each port and the food tray, whereas a house light was placed above the food tray. The right-hand side port of each chamber was designated as “active”, meaning that spontaneous nosepoking would result in the drop of one 45 mg food pellet (Bio-Serv, Flemington, NJ, USA) to the food tray. Upon nosepoking in the active port, the cue and house lights turned off, while the tray light turned on and the pellet dropped to the food tray. The two ports remained inactive for 20 s, during which nosepokes would not result in the delivery of a new pellet (time-out period). Subsequently, the chamber returned to its initial condition. Each training session lasted maximally two hours or until a rat acquired 100 pellets. Each rat received six training sessions (one training on each day for five consecutive days, followed by two days without training and one more training session on day 8). Only rats that finished at least two training sessions acquiring 100 pellets before the two-hour mark were used for progressive ratio reinforcement schedule (progressive ratio test) experiments. To test motivated behavior, rats were exposed to a progressive ratio test. Progressive ratio test sessions were identical to training sessions except that the operant requirement in each trial (T) was the integer (rounded down) of the function 1.4^(T-1)^ starting at one nosepoke for the first three trials and increasing in subsequent trials, as has been previously described^78^. Correct nosepokes (i.e. nosepokes in the active port and outside the timeout period, thus resulting in food delivery), number of obtained sucrose pellets (rewards) and the last ratio completed (breakpoint) were calculated to evaluate behavioral performance^19,79^.

### High Performance Liquid Chromatography (HPLC) analysis of GSH levels in brain samples

Animals were decapitated and their brains were quickly removed, frozen in isopentane on dry ice, at a temperature between −50 and −40°C, and stored at −80°C until further processing. Coronal sections (200-μm thick) were punched to obtain the brain tissue of NuAc region as previously described^80^

For GSH measurements, brain samples were briefly sonicated in Eppendorf vials containing 100 μl of 0.5 M perchloric acid (PCA) + 100μM EDTA-2Na and centrifuged at 16,000 *g* for 10 min at 4°C. The supernatant was collected, filtered through 0.22μm filters (5,000 g for 30” at 4°C) and used for high performance liquid chromatography (HPLC) analysis. Levels of GSH were assessed by reverse-phase HPLC with electrochemical detection (HPLC-ECD stand-alone system, HTEC-500). Using a mobile phase, consisting of 27.59g/l sodium phosphate monobasic monohydrate (pH2.5), 20 mL/l methanol, 10 mg/l EDTA-2Na and 100mg/l SOS dissolved in Milli-Q water, the GSH was separated in a reversed phase separation column EICOMPACK SC-3ODS using a Gold electrode (ref WE-AU).

### *Ex vivo* electrophysiology

Rats were anesthetized with isoflurane and decapitated. The brain was quickly removed, and coronal slices (250 μm thick) containing the ventral striatum were cut using a vibrating tissue slicer (Campden Instruments) in oxygenated (95% O_2_ / 5% CO_2_) ice-cold modified artificial CSF (ACSF), containing (in mM): 105 sucrose, 65 NaCl, 25 NaHCO3, 2.5 KCl, 1.25 NaH_2_PO_4_, 7 MgCl_2_, 0.5 CaCl_2_, 25 glucose, 1.7 L(+)-ascorbic acid. Slices recovered for 1 h at 35°C in standard ACSF containing (in mM): 130 NaCl, 25 NaHCO_3_, 2.5 KCl, 1.25 NaH_2_PO_4_, 1.2 MgCl_2_, 2 CaCl_2_, 18 glucose, 1.7 L(+)-ascorbic acid, and complemented with 2 Napyruvate and 3 myo-inositol. In the recording chamber, slices were superfused with oxygenated standard ACSF. Medium spiny neurons (MSNs) in the NuAc core were patched in the whole-cell configuration with borosilicate pipettes (3-4 MΩ) filled with a KGluconate- or CsGluconate intracellular solution complemented with 0.1% biocytin.

Measurements of intrinsic excitability were conducted at nearly physiological temperature (30-32°C), with an intracellular solution containing (in mM): 130 KGluconate, 10 KCl, 10 HEPES, 10 phosphocreatine, 0.2 EGTA, 4 Mg-ATP, 0.2 Na-GTP (290-300 mOsm, pH 7.2-7.3). To elicit neuronal firing, MSNs were held at −70 mV with direct current injections in the current clamp configuration, using bridge compensation, and depolarization was provided by 5-s long current ramps of increasing magnitude (maximal current ranging from 50 to 300 pA). Recordings were conducted during the first 5 min after establishment of the wholecell condition. The rheobase (minimal current required to elicit spiking) and the firing threshold were measured as the level of current and voltage, respectively, that induced the first action potential in the ramp protocol. Input resistance (Ri) and cell capacitance (Cm) were evaluated from the passive response to a −10 mV hyperpolarizing step provided from a holding potential of −60 mV.

Miniature excitatory postsynaptic currents (mEPSCs) were recorded in MSNs voltage-clamped at −60 mV using pipettes (3-4 MΩ) filled with (in mM): 120 CsGluconate, 10 CsCl, 10 HEPES, 10 phosphocreatine, 5 EGTA, 4 Mg-ATP (290-300 mOsm, pH 7.2-7.3). Recordings were conducted at room temperature in the presence of the GABA_A_R blocker picrotoxin (0.1 mM) and the Na^+^ channel blocker Tetrodotoxin (0.001 mM). Synaptic currents were acquired for 5 min starting from >8 min after the establishment of the wholecell configuration, to ensure proper diffusion of the intracellular solution.

At the end of each recording, the patch pipette was gently retracted from the cell body to cause membrane re-sealing. Slices were fixated in 4% PFA overnight, then stored in phosphate-buffered saline (PBS) complemented with 30% sucrose until processing.

Electrophysiological data were acquired through a Digidata1550A digitizer. Signals were amplified through a Multiclamp700B amplifier (Molecular Devices), sampled at 20 kHz and filtered at 10 kHz using Clampex10 (Molecular Devices). Data were analysed using Clampfit10 (Molecular Devices). For detection of mEPSCs, traces were filtered at 1 kHz and analysed using the MiniAnalysis Program (Synaptosoft Inc., Decatur, USA) setting a detection threshold of twice the root mean square of the noise level. Detected events were verified by visual inspection. To construct cumulative frequency plots, the first 200 events recorded in each cell were considered.

### *Post hoc* identification of D1- and D2-MSNs

*Post hoc* in-situ hybridization (RNAscope® Multiplex Fluorescent Reagent Kit v2. Assay, Advanced Cell Diagnostics, Inc.) for *Drd1* and *Drd2* was performed after patch-clamp electrophysiology following the manufacturer’s instructions, with some modifications. In order to visualize biocytin-filled neurons, brain slices were washed (5 min at RT x 3 times) in PBS and incubated (overnight at 4°C) with streptavidin-Alexa 488 (ThermoFisher Scientific, S-11223, 1:500) in PBS-triton 0.05%. In the following day, slices were washed (5 min at RT x 3 times) in PBS, mounted in Superfrost Plus™ Adhesion Microscope Slides (Thermo Scientific), baked at 60°C for 30 min and post-fixed in pre-chilled 4% PFA in PBS (15 min at 4°C). Afterwards, slices were dehydrated (50% EtOH for 5 min, 70% EtOH for 5 min, twice with 100% EtOH for 5 min at RT) and air dried (5 min). Endogenous peroxidase activity was blocked by RNAscope® H_2_O_2_ treatment (10 min at RT), and sections were permeabilized by RNAscope® Target Retrieval (5 min at 60°C) and a RNAscope® Protease III (30 min at 40°C). Next, slices were hybridized with pre-warmed probe pairs to target *Drd1* and *Drd2* mRNA (channel C1, Rn-Drd2-C1 no.315641) by incubation for 2 h at 40°C. Signal was amplified by incubation with RNAscope® Multiplex FL v2 AMP1 (30 min at 40ºC), RNAscope® Multiplex FL v2 AMP2 (30 min at 40°C) and RNAscope® Multiplex FL v2 AMP3 (15 min at 40°C). Channel C1 signals was developed by incubation with RNAscope® Multiplex FL v2 HRP-C1, followed by incubation with Opal 570 (1:1500 diluted in RNAscope® Multiplex TSA Buffer) and finally RNAscope® Multiplex FL v2 HRP blocker. The same procedure was used to target *Drd1* mRNA (channel C2, Rn-Drd1a-C2 no.317031-C2). The channel C2 signals was developed by incubation with RNAscope® Multiplex FL v2 HRP-C2, followed by incubation with Opal 690.

Confocal images were acquired on a LSM 700 confocal microscope (Carl Zeiss) with 20X/0.8 NA air and 40X/1.3 NA oil-immersion objectives (Bioimaging and Optics Platform, BIOP, EPFL). Biocytin-positive cells (113 successfully recovered out of 141 recorded cells) were identified as D1-MSNs (61 out of 113) or D2-MSNs (49 out of 113) based on reactivity for Drd1 and Drd2 RNA. One single neuron was D1-D2 double-positive, and other two neurons were D1-D2 double-negative.

### NeuN and caspase 3 immunohistochemistry

For NeuN and cleaved caspase 3 analysis, free-floating coronal sections (10μm thick) were mounted and fixed for 30 min with 4% PFA in PBS at RT. Then, rinsed with PBS-containing 0.3% Triton-X-100 (PBS-T), followed by a blocking step of 1 h incubation in PBS containing 3% BSA. Primary antibodies against NeuN (Abcam, ab104224, clone: 1B7, lot: GR3215839-1, dilution 1:000) or rabbit anti-cleaved caspase 3 (Cell Signaling, 9661s, 1:200) were diluted in blocking solution and sections incubated overnight at 4°C with gentle shaking. Sections were then washed three times in PBS and incubated with secondary antibodies (donkey anti-rabbit Cy3, Jackson ImmunoResearch, 1:500 dilution) for 2 h at room temperature. After three rinses in PBS, sections were incubated for 5 min with 4,6-diamidino-2-phenylindole (DAPI; Sigma, D9542), washed again three times in PBS and then mounted in Vectashield (Vector Labs). NeuN and cleaved caspase 3 expression were assessed in the NuAc using a LSM 710 laser-scanning confocal microscope (Carl Zeiss) imaged using a 10X with a 1.0 digital zoom. Images were analyzed for number of NeuN positive cells or fluorescence intensity for caspase-3 activation using ImageJ (NIH, USA). At least two sections from each animal were measured and averaged to generate one value per hemisphere per animal for each drug infusion

### Statistical analyses

Detailed parameters from statistical tests are reported in figure legends and in Table S1 indicating the statistical test used, sample size (‘N’), as well as degree of freedom, F and P values.

For participants, data were analyzed with the *jamovi* software v1.6. The Spearman correlation coefficient was used throughout to keep comparisons between correlation coefficients consistent, even if not all variables violate normality assumptions (tested with a Shapiro-Wilk test). The effect of GSH on the number of successful 1 CHF trials at the second half of the experiment was tested using a pre-post analysis, by adding the number of successful 1 CHF trials in the first half of the experiment as a covariate in an ANCOVA model. Assumptions of normality and homogeneity of variances were tested with the Shapiro-Wilk and Levene’s tests respectively. All tests were two-tailed with a critical probability of p≤0.05.

For animal experiments, statistical analyses were performed with Prism software v8.4.3 (GraphPad software, San Diego, CA, USA) or statistical package for social sciences (SPSS) version 17 (SPSS Inc, Chicago, IL, US) using a critical probability of p≤0.05. All values are represented as mean ± SEM. For two-group comparisons, one- or two-tailed t-test or Mann-Whitney test were applied, as appropriate. When the effect of more than one independent variable was analyzed, two- or three-way ANOVA was used, followed by post-hoc tests, as appropriate (Holm-Sidak post-hoc test). For survival analysis, the Mantel-Cox test was used.

## Acknowledgements

The authors thank Dr. Ting Yin and Dr. Hongxia Lei for expert help with the performance of MRS in rats. This project has been supported by grants from the Swiss National Science Foundation [CR20I3-146431, 31003A-176206 and −197942 to C.S.; and SNSF-Spark grant (No 196558) to J.R., European Union’s Horizon 2020 research and innovation programme under the Marie Sklodowska-Curie (N° 895562) to E.R-F, NCCR Synapsy (51NF40-158776 and 185897)], Société des Produits Nestlé SA; and intramural funding from the EPFL to C.S. FH is currently supported by a VISN7 research development award from the VA. The funding sources had no additional role in study design, in the collection, analysis and interpretation of data, in the writing of the report or in the decision to submit the paper for publication. This paper reflects only the authors’ views and the European Union is not liable for any use that may be made of the information contained therein. ^1^H-MRS experiments were also supported financially by the Center for Biomedical Imaging (CIBM) of the University of Lausanne (UNIL), University of Geneva (UNIGE), Geneva University Hospital (HUG), Lausanne University Hospital (CHUV), Swiss Federal Institute of Technology (EPFL) and the Leenaards and Louis-Jeantet Foundations.

## Competing interests

The authors declare not potential conflicts of interest with exception that L Trovò, P Steiner, and N Preitner are employees from Nestle S.A. and the work included in this manuscript was partially supported by a grant from Nestle S.A.

## Author contributions

C.S. and P.S. conceived the project with input from I.Z., F.H., L.T., A.S. and N.P.; I.Z., E.R-F., O.Z. and F.H. performed the behavioral and molecular experiments in rats with input from L.T.; E.R-F run the MRS in rats, and I.Z. and E.R-F., analyzed the data; S.A. performed and analyzed the *ex vivo* electrophysiological recordings; A.S, and L.X performed the behavioral and MRS experiments in human and J.R.; I.Z., E.R-F., F.H., S.A. and C.S. wrote the manuscript with input from all the authors; C.S. supervised and financially supported the project.

## Data availability

All data generated or analyzed during this study are included in the manuscript and supporting files.

## Notes

### Competing Interest Statement

The authors declare not potential conflicts of interest with exception that L Trovo, P Steiner, and N Preitner are employees from Nestle S.A. and the work included in this manuscript was partially supported by a grant from Nestle S.A.

## References

1 Chong, T. T. J., Bonnelle, V. & Husain, M. in Progress in Brain Research Vol. 229 (eds Bettina Studer & Stefan Knecht) 71–100 (Elsevier, 2016).

2 Lepine, J. A., Podsakoff, N. P. & Lepine, M. A. A Meta-Analytic Test of the Challenge Stressor–Hindrance Stressor Framework: An Explanation for Inconsistent Relationships Among Stressors and Performance. Academy of Management Journal 48, 764–775, doi:10.5465/amj.2005.18803921 (2005).

3 Duckworth, A. L., Eichstaedt, J. C. & Ungar, L. H. The Mechanics of Human Achievement. Soc Personal Psychol Compass 9, 359–369, doi:10.1111/spc3.12178 (2015).

4 Kanfer, R., Frese, M. & Johnson, R. E. Motivation related to work: A century of progress. Journal of Applied Psychology 102, 338–355, doi:10.1037/apl0000133 (2017).

5 Knutson, B., Taylor, J., Kaufman, M., Peterson, R. & Glover, G. Distributed Neural Representation of Expected Value. The Journal of Neuroscience 25, 4806, doi:10.1523/jneurosci.0642-05.2005 (2005).

6 Berchio, C., Rodrigues, J., Strasser, A., Michel, C. M. & Sandi, C. Trait anxiety on effort allocation to monetary incentives: a behavioral and high-density EEG study. Translational Psychiatry 9, 174, doi:10.1038/s41398-019-0508-4 (2019).

7 Epstein, J. et al. Lack of ventral striatal response to positive stimuli in depressed versus normal subjects. The American journal of psychiatry 163, 1784–1790, doi:10.1176/ajp.2006.163.10.1784 (2006).

8 Zald, D. H. & Treadway, M. T. Reward Processing, Neuroeconomics, and Psychopathology. Annual Review of Clinical Psychology 13, 471–495, doi:10.1146/annurev-clinpsy-032816-044957 (2017).

9 Pessiglione, M., Vinckier, F., Bouret, S., Daunizeau, J. & Le Bouc, R. Why not try harder? Computational approach to motivation deficits in neuro-psychiatric diseases. Brain 141, 629–650, doi:10.1093/brain/awx278 (2018).

10 Salamone, J. D., Yohn, S. E., López-Cruz, L., San Miguel, N. & Correa, M. Activational and effort-related aspects of motivation: neural mechanisms and implications for psychopathology. Brain : a journal of neurology 139, 1325–1347, doi:10.1093/brain/aww050 (2016).

11 Salamone, J. D., Correa, M., Farrar, A. & Mingote, S. M. Effort-related functions of nucleus accumbens dopamine and associated forebrain circuits. Psychopharmacology 191, 461–482, doi:10.1007/s00213-006-0668-9 (2007).

12 Croxson, P. L., Walton, M. E., Reilly, J. X., Behrens, T. E. J. & Rushworth, M. F. S. Effort-Based Cost–Benefit Valuation and the Human Brain. The Journal of Neuroscience 29, 4531, doi:10.1523/jneurosci.4515-08.2009 (2009).

13 Haber, S. N. Neuroanatomy of Reward: A View from the Ventral Striatum.

14 Schmidt, L., Lebreton, M., Cléry-Melin, M.-L., Daunizeau, J. & Pessiglione, M. Neural Mechanisms Underlying Motivation of Mental Versus Physical Effort. PLOS Biology 10, e1001266, doi:10.1371/journal.pbio.1001266 (2012).

15 Robinson, T. E., Yager, L. M., Cogan, E. S. & Saunders, B. T. On the motivational properties of reward cues: Individual differences. Neuropharmacology 76, 450–459, doi:https://doi.org/10.1016/j.neuropharm.2013.05.040 (2014).

16 Knutson, B., Adams, C. M., Fong, G. W. & Hommer, D. Anticipation of Increasing Monetary Reward Selectively Recruits Nucleus Accumbens. The Journal of Neuroscience 21, RC159, doi:10.1523/JNEUROSCI.21-16-j0002.2001 (2001).

17 Pessiglione, M. et al. How the Brain Translates Money into Force: A Neuroimaging Study of Subliminal Motivation. Science 316, 904–906, doi:10.1126/science.1140459 (2007).

18 Hailwood, J. M. et al. Oxygen responses within the nucleus accumbens are associated with individual differences in effort exertion in rats. European Journal of Neuroscience 48, 2971–2987, doi:https://doi.org/10.1111/ejn.14150 (2018).

19 Soares-Cunha, C. et al. Activation of D2 dopamine receptor-expressing neurons in the nucleus accumbens increases motivation. Nature Communications 7, doi:10.1038/ncomms11829 (2016).

20 Soares-Cunha, C. et al. Nucleus Accumbens Microcircuit Underlying D2-MSN-Driven Increase in Motivation. eneuro 5, ENEURO.0386-0318.2018, doi:10.1523/eneuro.0386-18.2018 (2018).

21 Gallo, E. F. et al. Accumbens dopamine D2 receptors increase motivation by decreasing inhibitory transmission to the ventral pallidum. Nature Communications 9, doi:10.1038/s41467-018-03272-2 (2018).

22 Epstein, J. et al. Lack of Ventral Striatal Response to Positive Stimuli in Depressed Versus Normal Subjects. American Journal of Psychiatry 163, 1784–1790, doi:10.1176/ajp.2006.163.10.1784 (2006).

23 Hanson, J. L., Hariri, A. R. & Williamson, D. E. Blunted Ventral Striatum Development in Adolescence Reflects Emotional Neglect and Predicts Depressive Symptoms. Biol Psychiatry 78, 598–605, doi:10.1016/j.biopsych.2015.05.010 (2015).

24 Muir, J., Lopez, J. & Bagot, R. C. Wiring the depressed brain: optogenetic and chemogenetic circuit interrogation in animal models of depression. Neuropsychopharmacology 44, 1013–1026, doi:10.1038/s41386-018-0291-6 (2019).

25 Treadway, M. T. et al. Corticolimbic gating of emotion-driven punishment. Nature Neuroscience 17, 1270–1275, doi:10.1038/nn.3781 (2014).

26 Hollis, F. et al. Mitochondrial function in the brain links anxiety with social subordination. Proceedings of the National Academy of Sciences of the United States of America 112, 15486–15491, doi:10.1073/pnas.1512653112 (2015).

27 van der Kooij, M. A. et al. Diazepam actions in the VTA enhance social dominance and mitochondrial function in the nucleus accumbens by activation of dopamine D1 receptors. Mol Psychiatry 23, 569–578, doi:10.1038/mp.2017.135 (2018).

28 Strasser, A. et al. Glutamine-to-glutamate ratio in the nucleus accumbens predicts effort-based motivated performance in humans. Neuropsychopharmacology 45, 2048–2057, doi:10.1038/s41386-020-0760-6 (2020).

29 Gebara, E. et al. Mitofusin-2 in the Nucleus Accumbens Regulates Anxiety and Depression-like Behaviors Through Mitochondrial and Neuronal Actions. Biol Psychiatry 89, 1033–1044 (2021).

30 Larrieu, T. et al. Hierarchical Status Predicts Behavioral Vulnerability and Nucleus Accumbens Metabolic Profile Following Chronic Social Defeat Stress. Current Biology 27, 2202–2210.e2204, doi:https://doi.org/10.1016/j.cub.2017.06.027 (2017).

31 Cherix, A. et al. Metabolic signature in nucleus accumbens for anti-depressant-like effects of acetyl-L-carnitine. eLife 9, e50631, doi:10.7554/eLife.50631 (2020).

32 Weger, M. et al. Mitochondrial gene signature in the prefrontal cortex for differential susceptibility to chronic stress. Scientific Reports 10, 18308, doi:10.1038/s41598-020-75326-9 (2020).

33 Hyder, F., Rothman, D. L. & Shulman, R. G. Total neuroenergetics support localized brain activity: Implications for the interpretation of fMRI. Proceedings of the National Academy of Sciences 99, 10771, doi:10.1073/pnas.132272299 (2002).

34 Smith, A. J. et al. Cerebral energetics and spiking frequency: The neurophysiological basis of fMRI. Proceedings of the National Academy of Sciences 99, 10765, doi:10.1073/pnas.132272199 (2002).

35 Walls, A. B., Waagepetersen, H. S., Bak, L. K., Schousboe, A. & Sonnewald, U. The Glutamine–Glutamate/GABA Cycle: Function, Regional Differences in Glutamate and GABA Production and Effects of Interference with GABA Metabolism. Neurochemical Research 40, 402–409, doi:10.1007/s11064-014-1473-1 (2015).

36 Forman, H. J., Zhang, H. & Rinna, A. Glutathione: Overview of its protective roles, measurement, and biosynthesis. Molecular Aspects of Medicine 30, 1–12, doi:https://doi.org/10.1016/j.mam.2008.08.006 (2009).

37 Sappington, D. R. et al. Glutamine drives glutathione synthesis and contributes to radiation sensitivity of A549 and H460 lung cancer cell lines. Biochimica et Biophysica Acta (BBA) - General Subjects 1860, 836–843, doi:https://doi.org/10.1016/j.bbagen.2016.01.021 (2016).

38 Rae, C. D. & Williams, S. R. Glutathione in the human brain: Review of its roles and measurement by magnetic resonance spectroscopy. Analytical Biochemistry 529, 127–143, doi:https://doi.org/10.1016/j.ab.2016.12.022 (2017).

39 Zalachoras, I. et al. Therapeutic potential of glutathione-enhancers in stress-related psychopathologies. Neuroscience & Biobehavioral Reviews 114, 134–155, doi:https://doi.org/10.1016/j.neubiorev.2020.03.015 (2020).

40 Cobley, J. N., Fiorello, M. L. & Bailey, D. M. 13 reasons why the brain is susceptible to oxidative stress. Redox Biol 15, 490–503, doi:10.1016/j.redox.2018.01.008 (2018).

41 Lapidus, K. A. B. et al. In vivo 1H MRS study of potential associations between glutathione, oxidative stress and anhedonia in major depressive disorder. Neuroscience Letters 569, 74–79, doi:https://doi.org/10.1016/j.neulet.2014.03.056 (2014).

42 Wang, A. M. et al. Assessing Brain Metabolism With 7-T Proton Magnetic Resonance Spectroscopy in Patients With First-Episode Psychosis. JAMA Psychiatry 76, 314–323, doi:10.1001/jamapsychiatry.2018.3637 (2019).

43 Proulx, C. D. et al. A neural pathway controlling motivation to exert effort. Proceedings of the National Academy of Sciences 115, 5792, doi:10.1073/pnas.1801837115 (2018).

44 Hodos, W. Progressive Ratio as a Measure of Reward Strength. Science 134, 943–944 (1961).

45 Stewart, W. J. Progressive reinforcement schedules: A review and evaluation. Australian Journal of Psychology 27, 9–22, doi:10.1080/00049537508255235 (1975).

46 Bottino, F. et al. In Vivo Brain GSH: MRS Methods and Clinical Applications. Antioxidants 10, 1407 (2021).

47 Floresco, S. B. The Nucleus Accumbens: An Interface Between Cognition, Emotion, and Action. Annual Review of Psychology 66, 25–52, doi:10.1146/annurev-psych-010213-115159 (2015).

48 Kravitz, A. V., Tye, L. D. & Kreitzer, A. C. Distinct roles for direct and indirect pathway striatal neurons in reinforcement. Nature Neuroscience 15, 816–818, doi:10.1038/nn.3100 (2012).

49 Lobo Mary, K. et al. Cell Type–Specific Loss of BDNF Signaling Mimics Optogenetic Control of Cocaine Reward. Science 330, 385–390, doi:10.1126/science.1188472 (2010).

50 Lin, Y., Stephenson, M. C., Xin, L., Napolitano, A. & Morris, P. G. Investigating the Metabolic Changes due to Visual Stimulation using Functional Proton Magnetic Resonance Spectroscopy at 7T. Journal of Cerebral Blood Flow & Metabolism 32, 1484–1495, doi:10.1038/jcbfm.2012.33 (2012).

51 Xin, L., Schaller, B., Mlynarik, V., Lu, H. & Gruetter, R. Proton T1 relaxation times of metabolites in human occipital white and gray matter at 7 T. Magnetic Resonance in Medicine 69, 931–936, doi:https://doi.org/10.1002/mrm.24352 (2013).

52 O’Doherty, J. et al. Dissociable Roles of Ventral and Dorsal Striatum in Instrumental Conditioning. Science 304, 452–454, doi:10.1126/science.1094285 (2004).

53 Yacubian, J. et al. Dissociable Systems for Gain- and Loss-Related Value Predictions and Errors of Prediction in the Human Brain. The Journal of Neuroscience 26, 9530, doi:10.1523/jneurosci.2915-06.2006 (2006).

54 Sescousse, G., Li, Y. & Dreher, J.-C. A common currency for the computation of motivational values in the human striatum. Social Cognitive and Affective Neuroscience 10, 467–473, doi:10.1093/scan/nsu074 (2015).

55 Hall, C. N., Klein-Flügge, M. C., Howarth, C. & Attwell, D. Oxidative Phosphorylation, Not Glycolysis, Powers Presynaptic and Postsynaptic Mechanisms Underlying Brain Information Processing. The Journal of Neuroscience 32, 8940, doi:10.1523/jneurosci.0026-12.2012 (2012).

56 Schneider, J. et al. Local oxygen homeostasis during various neuronal network activity states in the mouse hippocampus. Journal of Cerebral Blood Flow & Metabolism 39, 859–873, doi:10.1177/0271678x17740091 (2019).

57 Dringen, R. Metabolism and functions of glutathione in brain. Progress in Neurobiology 62, 649–671, doi:https://doi.org/10.1016/S0301-0082(99)00060-X (2000).

58 Fernandez-Fernandez, S. et al. Hippocampal neurons require a large pool of glutathione to sustain dendrite integrity and cognitive function. Redox Biol 19, 52–61, doi:10.1016/j.redox.2018.08.003 (2018).

59 Sedlak, T. W. et al. The glutathione cycle shapes synaptic glutamate activity. Proceedings of the National Academy of Sciences 116, 2701–2706, doi:10.1073/pnas.1817885116 (2019).

60 Stuber Garret, D. et al. Reward-Predictive Cues Enhance Excitatory Synaptic Strength onto Midbrain Dopamine Neurons. Science 321, 1690–1692, doi:10.1126/science.1160873 (2008).

61 Day, J. J., Roitman, M. F., Wightman, R. M. & Carelli, R. M. Associative learning mediates dynamic shifts in dopamine signaling in the nucleus accumbens. Nature Neuroscience 10, 1020–1028, doi:10.1038/nn1923 (2007).

62 Flagel, S. B. et al. A selective role for dopamine in stimulus-reward learning. Nature 469, 53–57, doi:10.1038/nature09588 (2011).

63 Ducrocq, F. et al. Causal Link between n-3 Polyunsaturated Fatty Acid Deficiency and Motivation Deficits. Cell Metabolism 31, 755–772.e757, doi:https://doi.org/10.1016/j.cmet.2020.02.012 (2020).

64 Trifilieff, P. et al. Increasing dopamine D2 receptor expression in the adult nucleus accumbens enhances motivation. Molecular Psychiatry 18, 1025–1033, doi:10.1038/mp.2013.57 (2013).

65 Filla, I. et al. Striatal dopamine D2 receptors regulate effort but not value-based decision making and alter the dopaminergic encoding of cost. Neuropsychopharmacology 43, 2180–2189, doi:10.1038/s41386-018-0159-9 (2018).

66 Soares-Cunha, C. et al. Nucleus accumbens medium spiny neurons subtypes signal both reward and aversion. Molecular Psychiatry 25, 3241–3255, doi:10.1038/s41380-019-0484-3 (2020).

67 Tardiolo, G., Bramanti, P. & Mazzon, E. Overview on the Effects of N-Acetylcysteine in Neurodegenerative Diseases. Molecules 23, doi:10.3390/molecules23123305 (2018).

68 Koga, M. et al. Glutathione is a physiologic reservoir of neuronal glutamate. Biochemical and Biophysical Research Communications 409, 596–602, doi:https://doi.org/10.1016/j.bbrc.2011.04.087 (2011).

69 Zinsmaier, A. K., Dong, Y. & Huang, Y. H. Cocaine-induced projection-specific and cell type-specific adaptations in the nucleus accumbens. Molecular Psychiatry, doi:10.1038/s41380-021-01112-2 (2021).

70 Olive, M. F., Cleva, R. M., Kalivas, P. W. & Malcolm, R. J. Glutamatergic medications for the treatment of drug and behavioral addictions. Pharmacology Biochemistry and Behavior 100, 801–810, doi:https://doi.org/10.1016/j.pbb.2011.04.015 (2012).

71 van ‘t Erve, T. J., Wagner, B. A., Ryckman, K. K., Raife, T. J. & Buettner, G. R. The concentration of glutathione in human erythrocytes is a heritable trait. Free Radical Biology and Medicine 65, 742–749, doi:https://doi.org/10.1016/j.freeradbiomed.2013.08.002 (2013).

72 Dasari, S. et al. Genetic polymorphism of glutathione S-transferases: Relevance to neurological disorders. Pathophysiology 25, 285–292, doi:https://doi.org/10.1016/j.pathophys.2018.06.001 (2018).

73 Sohal, R. S., Ku, H.-H., Agarwal, S., Forster, M. J. & Lal, H. Oxidative damage, mitochondrial oxidant generation and antioxidant defenses during aging and in response to food restriction in the mouse. Mechanisms of Ageing and Development 74, 121–133, doi:https://doi.org/10.1016/0047-6374(94)90104-X (1994).

74 Xin, L. et al. Genetic Polymorphism Associated Prefrontal Glutathione and Its Coupling With Brain Glutamate and Peripheral Redox Status in Early Psychosis. Schizophr Bull 42, 1185–1196, doi:10.1093/schbul/sbw038 (2016).

75 Herrero, A. I., Sandi, C. & Venero, C. Individual differences in anxiety trait are related to spatial learning abilities and hippocampal expression of mineralocorticoid receptors. Neurobiology of Learning and Memory 86, 150–159, doi:https://doi.org/10.1016/j.nlm.2006.02.001 (2006).

76 van der Kooij, M. A. et al. Diazepam actions in the VTA enhance social dominance and mitochondrial function in the nucleus accumbens by activation of dopamine D1 receptors. Molecular Psychiatry 23, 569–578, doi:10.1038/mp.2017.135 (2017).

77 van der Kooij, M. A., Zalachoras, I. & Sandi, C. GABAA receptors in the ventral tegmental area control the outcome of a social competition in rats. Neuropharmacology 138, 275–281, doi:https://doi.org/10.1016/j.neuropharm.2018.06.023 (2018).

78 Wanat, M. J., Bonci, A. & Phillips, P. E. M. CRF acts in the midbrain to attenuate accumbens dopamine release to rewards but not their predictors. Nat Neurosci 16, 383–385, doi:http://www.nature.com/neuro/journal/v16/n4/abs/nn.3335.html#supplementary-information (2013).

79 Richardson, N. R. & Roberts, D. C. S. Progressive ratio schedules in drug self-administration studies in rats: a method to evaluate reinforcing efficacy. Journal of Neuroscience Methods 66, 1–11, doi:https://doi.org/10.1016/0165-0270(95)00153-0 (1996).

80 Guitart, X. et al. Regulation of Ionotropic Glutamate Receptor Subunits in Different Rat Brain Areas by a Preferential Sigma1 Receptor Ligand and Potential Atypical Antipsychotic. Neuropsychopharmacology 23, 539–546, doi:10.1016/s0893-133x(00)00142-1 (2000).

