## Supplementary Information for "Glutathione in the nucleus accumbens regulates motivation to exert reward-incentivized effort"

**Table of contents**

Supplementary Figure 1. GSH levels in the nucleus accumbens are not associated with the number of successful trials for lower rewards (0.5 CHF and 0.2 CHF).

Supplementary Figure 2. High accumbal GSH levels affects the probability to get a higher breakpoint and the persistence of nose poking over time.

Supplementary Figure 3. Accumbal BSO infusion decreases the probability of reaching higher breakpoints in PR task.

Supplementary Figure 4. NAC treatment affects the probability of obtaining a higher breakpoint and persistence of nose poking over the time but does not change body weight or exploration levels.

Supplementary Figure 5. NAC treatment does not affect MSN intrinsic excitability.

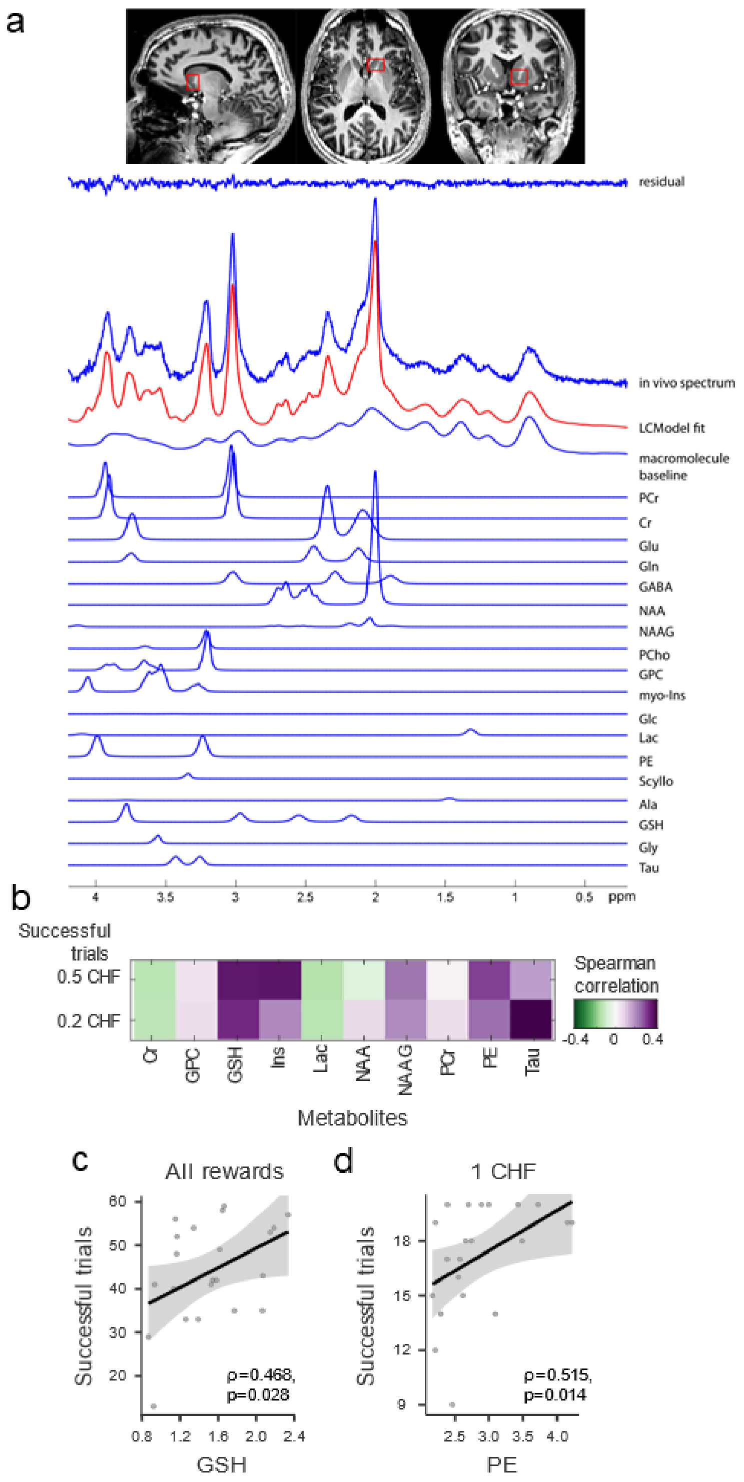


**Supplementary Figure 1. GSH levels in the nucleus accumbens are not associated with the number of successful trials for lower rewards (0.5 CHF and 0.2 CHF).**

(**a**) Region of interest in the human brain (i.e., nucleus accumbens) and examples of metabolites spectra. (**b**) Matrix showing all pairwise correlations between metabolites of interest and the number of successful trials in the entire task for 0.5 CHF and 0.2 CHF rewards (**c**). Detailed scatterplot for the correlation between GSH levels in the nucleus accumbens and the number of successful trials for all monetary rewards (ρ=0.468, p=0.028). (**d**) Detailed scatterplot for the correlation between PE levels in the nucleus accumbens and the number of successful 1 CHF trials (ρ=0.515, p=0.014).

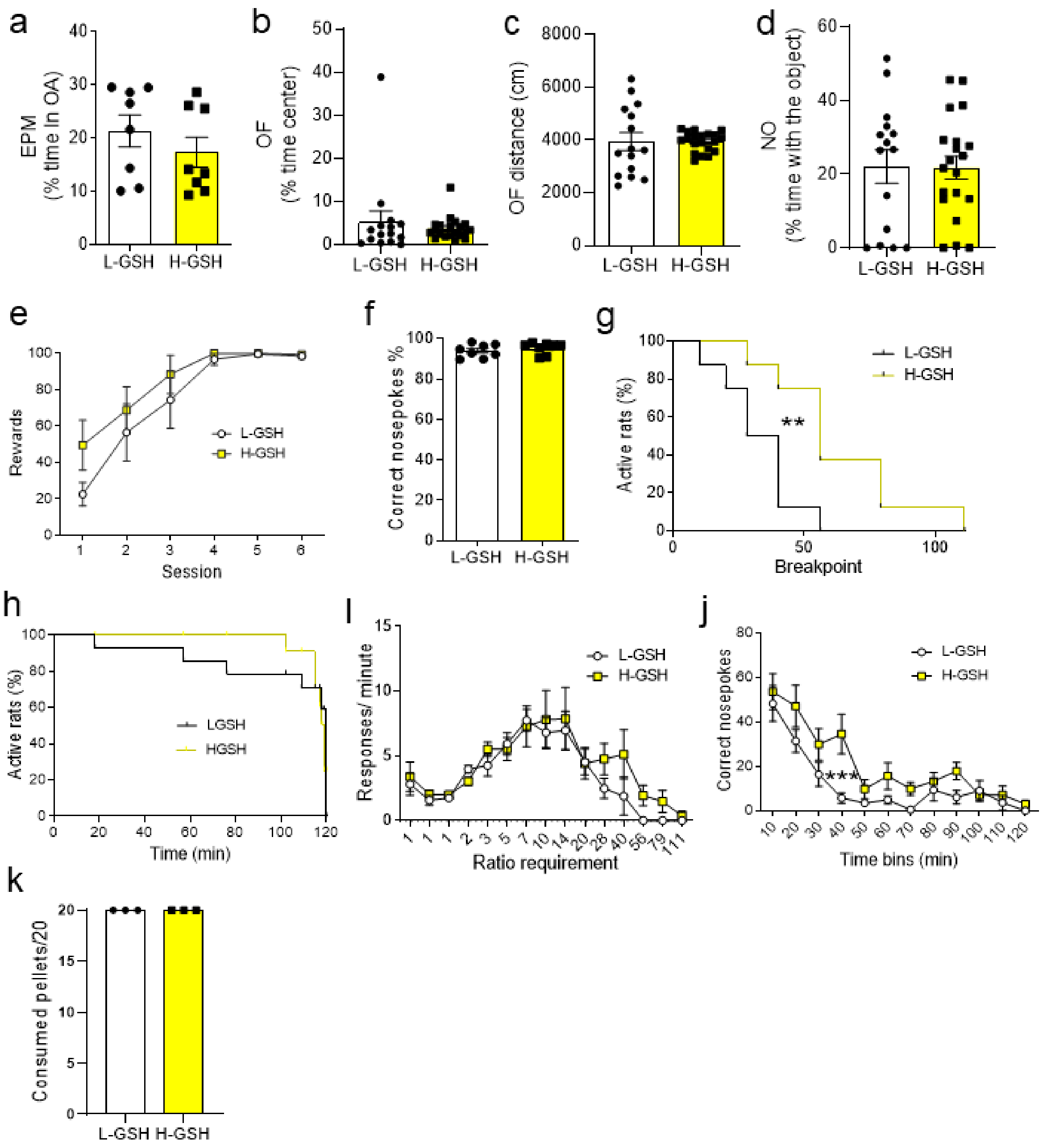


**Supplementary Figure 2. High accumbal GSH levels affects the probability to get a higher breakpoint and the persistence of nose poking over time.**

(**a**) High- (H) and Low (L)-GSH rats showed no differences in the percentage of time spent in the open arms of the EPM (t(14) = 0.9686, p = 0.3492). No difference was observed between the two groups in the time the rats spent in the center of the open field (t(33) = 0.7137, p = 0.4804), (**b**) in the distance travelled in the open field (t(33) = 0.0137, p = 0.9891) or (**c**) the percentage of time spent exploring a novel object (**d**) (t(33) = 0.04682, p = 0.9629). (**e**)There were no differences between H-GSH and L-GSH rats in the training performance in the operant conditioning. Two-way repeated measures ANOVA revealed a non-significant Time X GSH interaction (F(5,65) = 0.7966, p = 0.5561), a significant effect of time (F(5,65) = 20.04, p < 0.0001), and a non-significant effect of GSH (F(1,13) = 1.615, p = 0.226). (**f**) The percentage of correct nosepokes over total nosepokes in the PR test was not different between the groups ($t(14) = 0.8828$, $p = 0.3923$). (**g**) Survival analysis showed that the H-GSH group had higher probability to reach higher breakpoints than the L-GSH group (Long-rank Mantel-Cox test, Chi2 = 6.879, $p < 0.01$). (**h**) Survival analysis showed that L-GSH and H-GSH rats did not show differences in how long they kept performing nosepokes (Long-rank Mantel-Cox test, Chi2 = 0.0, $p = 0.999$). (**i**)There were no differences between groups considering the response/minute by ratio requirements. Two-way repeated measures ANOVA showed a non-significant GSH x ratio requirements (F14,168) = 0.737, $p = 0.735$), a significant effect of ratio-requirement ($F(14, 168) = 13.87$, $p < 0.001$) and a non-significant effect of GSH ($F(1,12) = 0.899$, $p = 0.362$). (**j**) When considering the number of correct nosepokes over time, two-way repeated measures ANOVA revealed a non-significant GSH X Time interaction ($F(11, 154) = 1.534$, $p = 0.1246$) a significant effect of time ($F(11,154) = 21.85$, $p < 0.001$) and a significant effect of GSH levels ($F(1,14) = 6.811$, $p = 0.0206$). (**k**) L-GSH and H_GSH rats did not exhibit differences in the consumption of pellets when they were given free access to them.

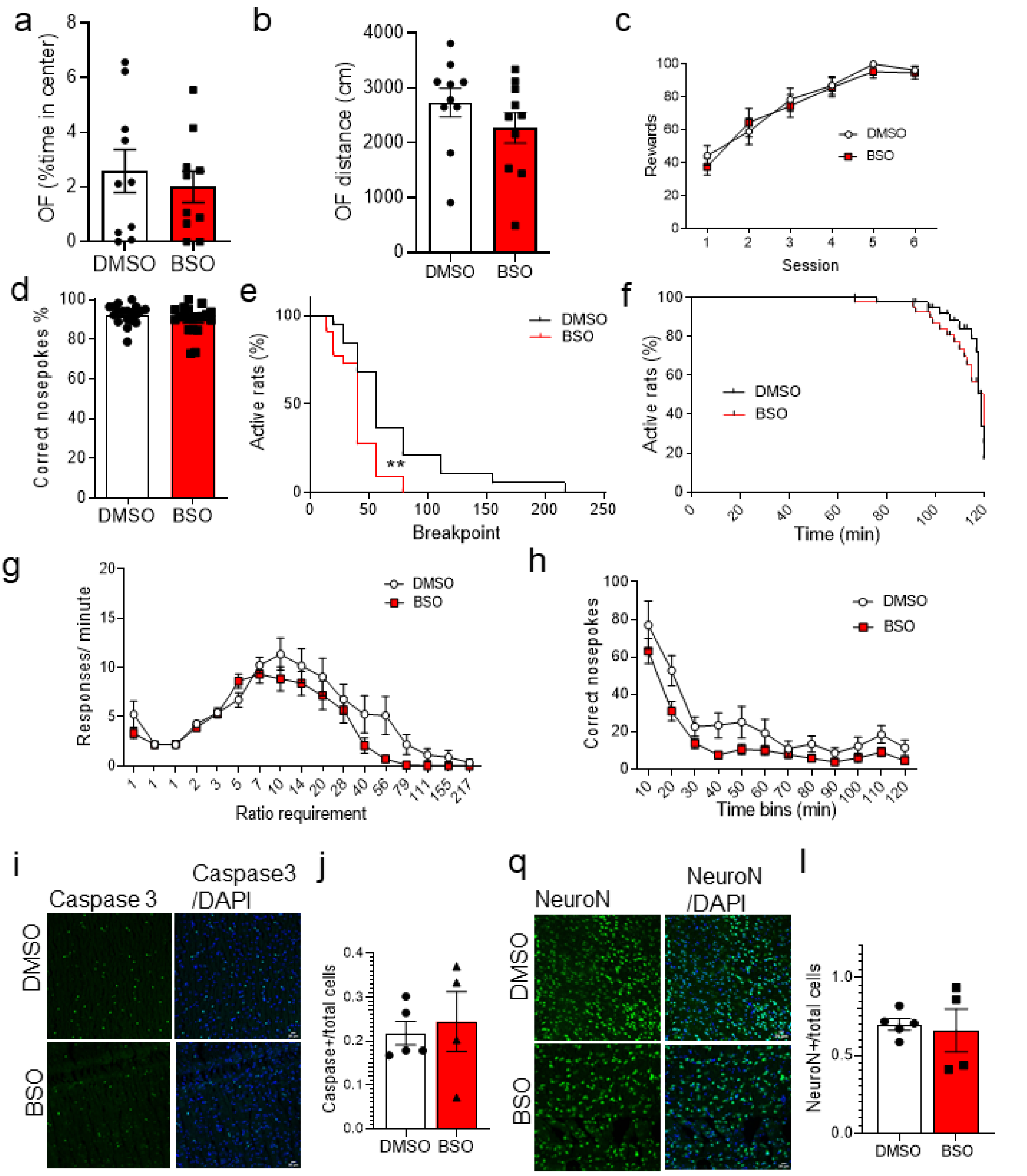


**Supplementary Figure 3. Accumbal BSO infusion decreases the probability of reaching higher breakpoints in PR task.**

(**a**) No differences were observed in the percentage of time spent in the center of the open field (unpaired two-tailed t-test, t(18) = 0.5959, p = 0.5586) or (**b**) the distance travelled in the open field between groups (unpaired two-tailed t-test, t(18) = 1.203, p = 0.2445). (**c**) The DMSO and BSO groups did not show differences for their operant training performance in the FR1 training. Repeated measures two-way ANOVA showed a non-significant group x time interaction ($F_{(5, 230)}$ = 0.4134, p = 0.8392), a significant effect of time (F(5, 230) = 45.27, p < 0.0001) and a non-significant effect of treatment (F(1, 46) = 0.144, p = 0.7060). (**d**) Regarding the percentage of correct nosepokes over the total number of nosepokes performed in the PR test, there were no differences between groups (unpaired two-tailed t-test, t(37) = 1.343, p = 0.1875). (**e**) There was a lower probability of reaching a higher breakpoint for the BSO group than the DMSO (Long-rank Mantel-Cox test, Chi2(1) = 7.874, p = 0.005). (**f**) BSO rats did not show a significant decrease in the probability to make correct nosepokes over time, compared to the saline group (Long-rank Mantel-Cox test, Chi2 (1) = 0.2660, p = 0.6061). (**g**) No difference was observed in the rate of responses per ratio requirement between the two groups. Two-way ANOVA revealed a non-significant ratio requirement X treatment interaction (F(16, 608) = 1.494, p = 0.0961), a significant ratio requirement effect (F(16, 608) = 32.93, p < 0.0001) and a non-significant effect of treatment (F(1,38) = 2.605, p = 0.1148). (**h**) There was a non-significant treatment x time interaction (F(11, 418) = 0.7685, p = 0.6717), but significant effects of time (F(11, 418) = 36.47, p < 0.0001) and treatment (F(1, 38) = 6.172, p = 0.0175) when comparing the correct nosepokes over time. (**i** and **j**) Confocal images and the quantification showing no differences in Caspase 3 activation following BSO treatment in the nucleus accumbens (unpaired two-tailed t-test, t(7) = 0.3905, p = 0.7078). (**q** and **l**) NeuN staining revealed no difference in the number of NeuN + cells in the nucleus accumbens, following BSO treatment (unpaired two-tailed t-test, t(7) = 0.3098, p = 0.7657). N=4-5/group.

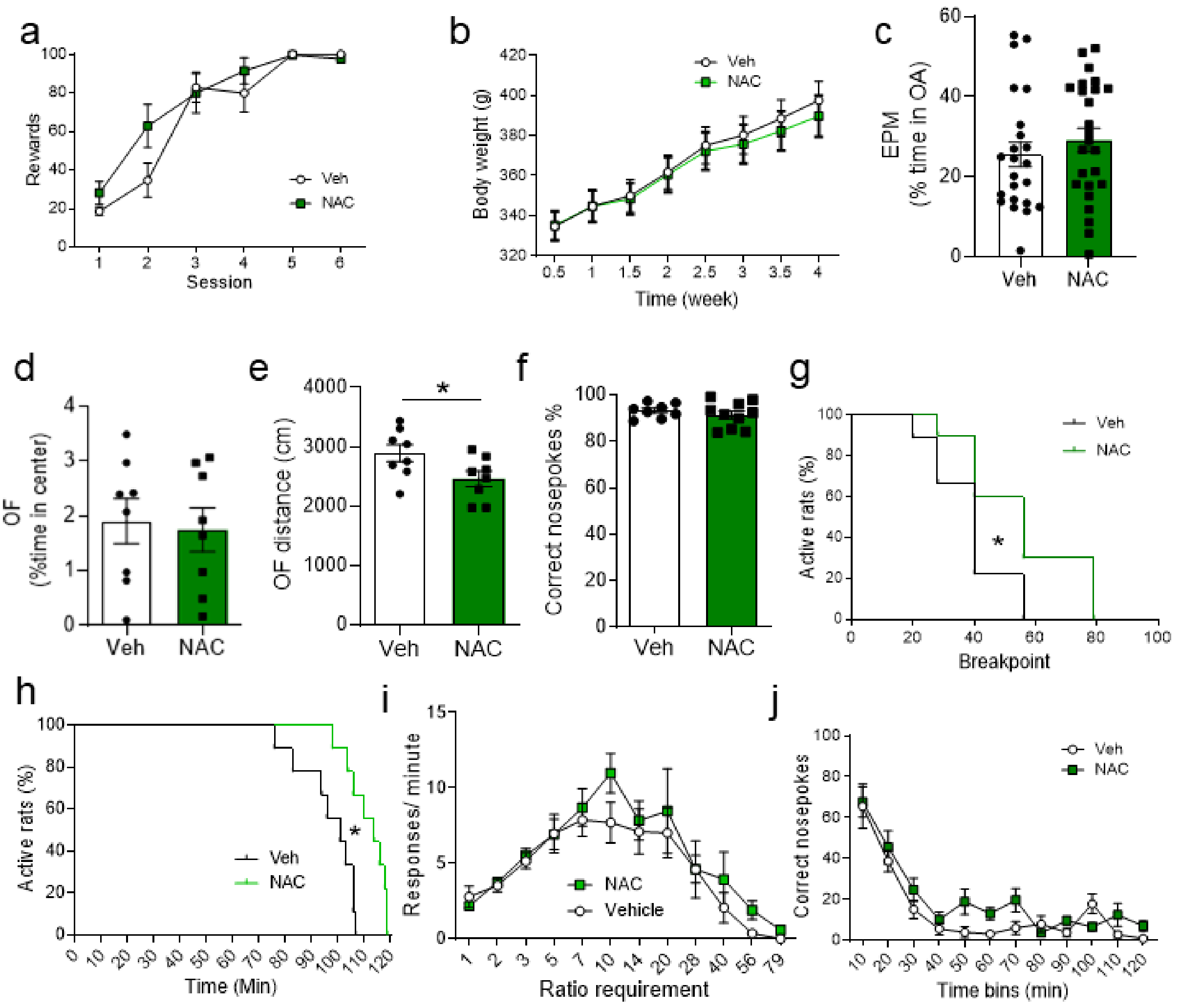

**Supplementary Figure 4. NAC treatment affects the probability of obtaining a higher breakpoint and persistence of nose poking over the time but does not change body weight or exploration levels.**
(**a**) During operant conditioning training, no differences were observed between Veh and NAC treated groups (Two-way repeated measures ANOVA, Treatment: F(1,19) = 1.272, p = 0.2734, Time: F(5,95) = 59.32, p < 0.001, Time X treatment interaction: F(5,95) = 2.263, p = 0.0543). (**b**) The two groups did not differ in body weight over time in animals neither at baseline or after treatment (Two-way repeated measures ANOVA, Treatment: F(1,20) = 0.057, p = 0.8146, Time: F(7,140) = 449.9, p < 0.001, Time X treatment interaction: F(7, 140) = 2.293, p = 0.0305), nor in the percent time spent in EPM the open arm before the treatment (unpaired two-tailed t-test, t(46) = 0.8183, p = 0.4174) (**c**). (**d**) NAC-treated rats from a different cohort showed no difference in the percentage of time spent in the center of the open field compared to the Veh group (unpaired two-tailed t-test, t(14) = 0.2793, p = 0.7841). (**e**) However, rats that received NAC treatment travelled a shorter distance in the open field test (unpaired two-tailed t-test, t(14) = 2.224, p = 0.0431). (**f**) During the PR task, no differences were observed between Veh- and NAC-treated groups on the percentage of correct nosepokes over the total number of nosepokes (unpaired two-tailed t-test, t(16) = 0.8225, p = 0.4229). (**g**) Interestingly, the survival plots showed that NAC-treated rats had a higher probability to reach higher breakpoints than Veh-treated rats Long-rank (Mantel-Cox test, Chi2(1) = 4.362, p = 0.0368). (**h**) NAC-treated rats had a higher probability to continue performing nosepokes for longer (Long-rank Mantel-Cox test, Chi2(1) = 9.278, p = 0.0023). (**i**) No statistically significant differences were found between groups regarding the

number of responses/minute for the different ratio requirements (Two-way repeated measures ANOVA, Treatment: $F(1,17) = 0.9195$, $p = 0.3510$, Time: $F(11,187) = 15.45$, $p < 0.001$, Time X treatment interaction: $F(11,187) = 0.4885$, $p = 0.9089$), or the correct nosepokes over time (**j**) (Two-way repeated measures ANOVA, Treatment: $F(1,17) = 3.717$, $p = 0.0707$, Time: $F(11,187) = 34.93$, $p < 0.001$, Time X treatment interaction: $F(11,187) = 1.381$, $p = 0.1848$).

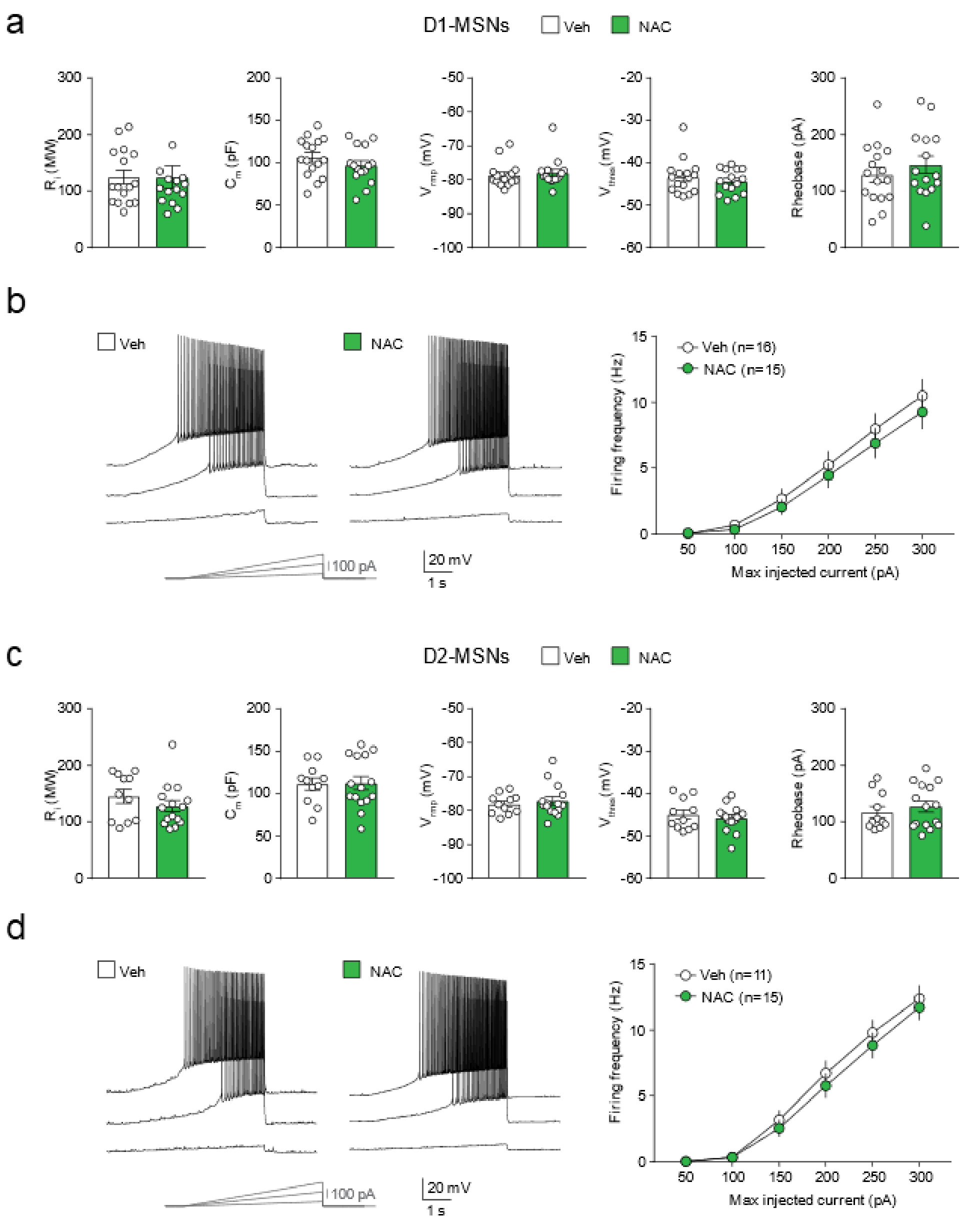


**Supplementary Figure 5. NAC treatment does not affect MSN intrinsic excitability.**

(**a**) Lack of differences in cell passive and active properties of NAc core D1-MSNs from vehicle- and NAC-treated rats: input resistance ($R_i$) (Mann-Whitney test, U=110, p=0.7112); membrane capacitance ($C_m$) (unpaired t test, t=1.167, p=0.2526); resting membrane potential ($V_{rmp}$) (Mann-Whitney test, U=92, p=0.2814); firing threshold ($V_{thres}$) (Mann-Whitney test, U=111.5, p=0.7477); rheobase (unpaired t test, t=0.8658, p=0.394). (**b**) Left, example voltage responses elicited in D1-MSNs from vehicle- and NAC-treated rats by injection of current ramps (protocol at the bottom; 3 of 6 ramps displayed). Right, frequency-current relationship of action potential discharges evoked by current ramps, indicating no treatment-induced differences (Two-way ANOVA for factor treatment, $F_{(1, 174)}$=1.695, p=0.1946). (**c**) Lack of differences in cell passive and active properties of nucleus accumbens core D2-MSNs from vehicle- and NAC-treated rats: $R_i$ (Mann-Whitney test, U=61, p=0.2754), $C_m$ (unpaired t test, t=1.1084, p=0.9146); $V_{rmp}$ (Mann-Whitney test, U=78, p=0.8384); $V_{thres}$ (unpaired t test, t=0.639, p=0.5304); rheobase (Mann-Whitney test, U=73, p=0.6368). (**d**) Same representation as in **b** for D2-MSNs (Two-way ANOVA for factor treatment, $F_{(1, 144)}$=1.477, p=0.2262).
